# Rate limiting release of product underlies concave Arrhenius break point of thermolysin with a Phe-Leu-Ala substrate

**DOI:** 10.64898/2026.04.22.720203

**Authors:** Jared J. Miller, Brian J. Bahnson

## Abstract

Thermolysin, a bacterial zinc metalloprotease, has been previously been reported to exhibit a biphasic kinetic temperature dependence of *k*_*cat*_ with a characteristic convex shape. This convex shaping is observed for almost all enzymes which display an Arrhenius break; fumarase is the exception with concave shaping. Here, thermolysin kinetics measured with the tripeptide substrate N-[3-(2-furyl)acryloyl]-Phe-Leu-Ala (FAFLA) resulted in a concave Arrhenius plot, characterized by a 30 kJ/mol increase in enthalpy and entropy of activation, in contrast to the typical 30 kJ/mol decrease. Although the shape of the Arrhenius break differs, ionic strength and macromolecular crowding both attenuate the energetic magnitude of the break point, consistent with prior work. It was hypothesized that a different step of the catalytic cycle of thermolysin was represented by *k*_*cat*_ with FAFLA to give rise to this new behavior. A 91% dependence of *k*_*cat*_ on viscosity and modest solvent isotope effects, both distinct from previously-characterized substrates, indicated that a physical step was responsible for the observed Arrhenius concavity. Hinge bending conformational changes of thermolysin, monitored using the phosphoramidon inhibitor (a FAFLA mimic), exhibited a fully linear temperature dependence, excluding these large-scale motions as the origin of concavity. It was therefore proposed that release of the N-[3-(2-furyl)acryloyl]-Phe product is likely rate limiting since release was proposed to involve a two-step pathway to free the product coordinated to the catalytic Zn^2+^ of thermolysin. These findings provide a mechanistic framework for seldom-seen concave break point behavior and insights into the contribution of dynamics of physical processes to catalysis.

**IMPORTANCE AND IMPACT:** Enzymes which display Arrhenius break behavior provide insight into how dynamics impact catalysis. Almost every enzyme thus far displays convex biphasic shape, with concave shaping often not acknowledged. Thermolysin, which previously only showed convex shaping, displayed concave behavior with a tripeptide substrate. By linking this unusual kinetic behavior to a physical, not chemical, process, this work highlights the possible origin of a rare phenomenon which can expand understanding of protein dynamics and biphasic Arrhenius behavior.

## INTRODUCTION

The Arrhenius and Eyring equations model the typical, single phase linear relationship between temperature and reaction rate of an enzyme. However, some enzymes instead demonstrate biphasic kinetic temperature dependences: two linear regimes which meet at an Arrhenius break point which has been linked to a change in structural dynamics at this temperature^1–14^. Almost all enzymes with biphasic Arrhenius behavior display convex shaping (Figure 1), which is the result of a concurrent decrease in enthalpy (ΔH^‡^) and entropy (TΔS^‡^) of activation across the break point^1,15,16^. Concave break point behavior (an increase in ΔH^‡^ and TΔS^‡^ with increasing temperature) has only been consistently observed with fumarase^9,11^, although Tolman’s equation suggests that concave Arrhenius plots should be more common due to activation energy (E_a_) tending to increase with temperature^16^. Phosphorylase kinase (PhK) and acetylcholinesterase (AchE) have been reported to show concave behavior under some conditions, but these cases are ambiguous: the temperature dependence of PhK linearized upon study with a lower enzyme concentration^14^ and AchE showed an atypically large 120 kJ/mol change in activation parameters across the break point^13^.

**Figure 1.**
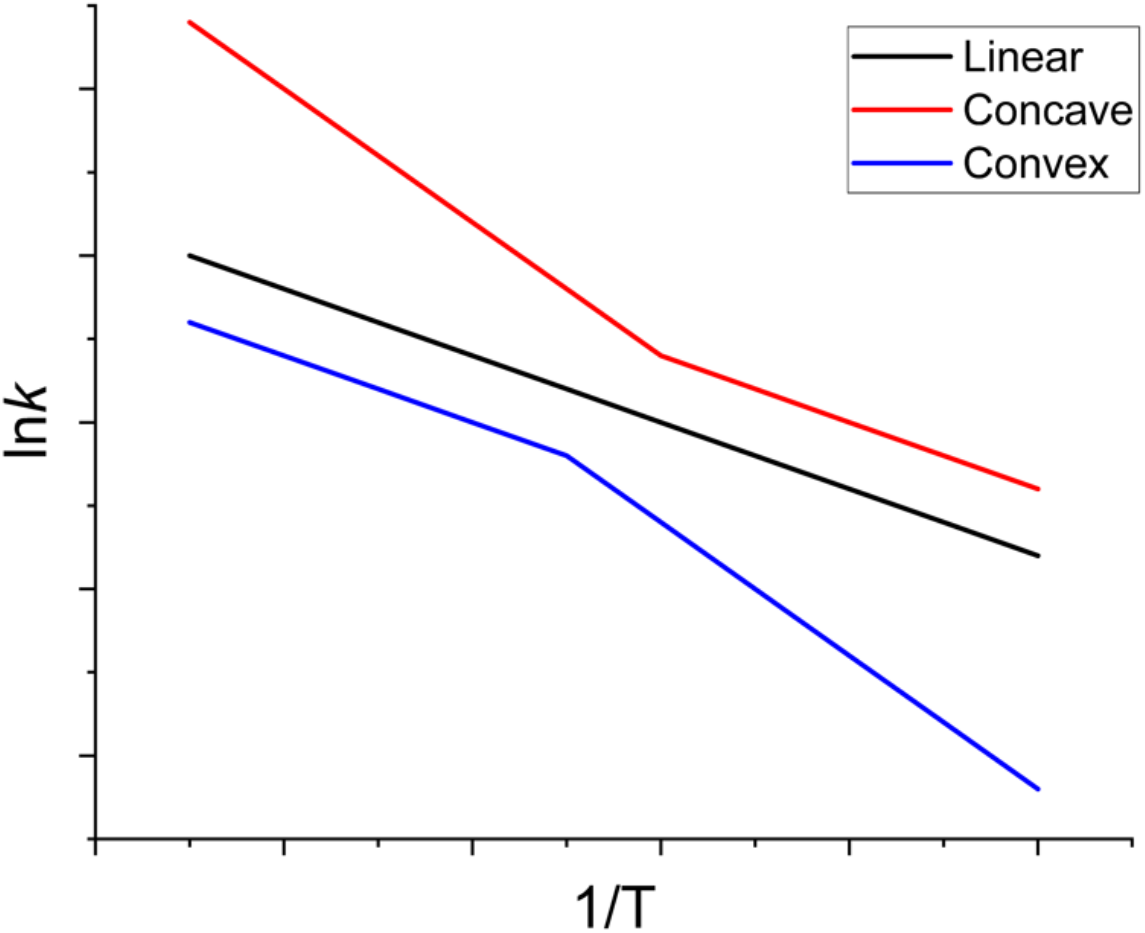
Types of Arrhenius Plots. Kinetic temperature dependences of enzymes can take on three shapes, with linear (black line) being standard behavior modeled by the Arrhenius and Eyring equations^16^. Convex (blue) biphasic Arrhenius plots represent the behavior of most enzymes characterized with biphasic Arrhenius behavior thus far. Concave (red) behavior has only been consistently observed with fumarase^9,11^.

Previously, the protease thermolysin demonstrated convex Arrhenius behavior with peptide and protein substrates^1,2,17–19^. Here, thermolysin displayed concave break point shaping with the tripeptide substrate N-[3-(2-furyl)acryloyl]-Phe-Leu-Ala (FAFLA), representing a novel example of concavity outside fumarase.

Phe-Leu-Ala (FLA) based ligands have previously demonstrated unique behavior with thermolysin that must be considered. The substrate N-benzyloxycarbonyl-Phe-Leu-Ala (ZFLA) previously demonstrated *k*_*cat*_/K_M_ values ranging from 700-1300 mM^−1^s^−1^ with k_cat_ values ranging between 300-700 s^−1^; both of which being uniquely large for thermolysin with peptide substrates^20,21^. The phosphoramide transition state analog N-benzyloxycarbonyl-Phe-phosphoryl-Leu-Ala (ZF^P^LA) is the most potent inhibitor of thermolysin with a K_i_ of 68 pM^22^, with this remarkable affinity found to be the result of steric interactions between the Phe residue of the substrate and Asn112 of thermolysin^23^. These steric interactions were proposed to stabilize the closed, ligand-bound form of the enzyme, dramatically increasing the residence time from 3 min with N-benzyloxycarbonyl-Gly-phosphoryl-Leu-Ala (ZG^P^LA) to 168 d with ZF^P^LA^23^. Furthermore, FLA ligands were found to bind in a distinct mode, which excluded additional water and was deeper into the active site cleft than other ligands^24^.

The focus of this study is to determine which step of the catalytic pathway of thermolysin is responsible for the concave break point shape with FAFLA. The convex break point of the chemical steps of thermolysin have been characterized previously with N-[3-(2-furyl)acryloyl]-Gly-Leu-Amide (FAGLA) and related analogs^2,17–19^, and a distinct physical step has been examined with the succinylated bovine β-casein (succinylcasein) substrate^1^. Since fumarase, AchE, and PK are not limited by chemical steps^25–27^, it was hypothesized that a physical step in the catalytic cycle of thermolysin would be underlie this concave behavior.

To test this, kinetic solvent viscosity (KSVE) and kinetic solvent isotope (KSIE) effects were used to determine if the rate-limiting step of FAFLA turnover is chemical or physical and whether this step is different between FAFLA and previously-characterized substrates. The hinge bending motion of thermolysin upon substrate binding (Figure S1)^24,28^ was examined using the FAFLA-mimic phosphoramide transition state analog phosphoramidon (Figure S2)^24,29^ to determine if this or the retrograde motion during product release are the origin of concave break point shape. Finally, the break point of thermolysin with the FAFLA substrate was examined in the presence of elevated NaCl (ionic strength) and Ficoll-70 (macromolecular crowding), which had previously been shown to diminish the energetic difference across the break point, to evaluate whether this effect persists with a different substrate and break point shape.

## RESULTS

### Break Point Behavior of Thermolysin with the FAFLA Substrate

The Arrhenius plot of thermolysin with the FAFLA substrate shows concave behavior with a break point temperature range of 24-28 °C among all experiments, which is consistent with previous characterizations of convex shaping (Figure 2A)^1,2,17–19^. The difference in ΔH^‡^ and TΔS^‡^ across the break point will be referred to as ΔH°_c_ and TΔS°_c_, respectively, which quantify the energetic change associated with the break point transition^30^. Instead of the typical 28 kJ/mol decrease^1^, the concave shaping now demonstrates a 34 ± 8 kJ/mol increase in ΔH°_c_ and TΔS°_c_ at the break point (Figure 2B). ΔG^‡^ increases with temperature, and the break point reduces the rate at which ΔG^‡^ increases at higher temperatures (Figure S3). The break point seems to continue to originate solely from *k*_*cat*_^1,17^, as *k*_*cat*_ values can be superimposed onto the Arrhenius plot collected at 0.70 mM FAFLA (Figure 2C). K_M_ values show no significant temperature dependence and do not contribute appreciably to the concave behavior (Figure 2D).

**Figure 2.**
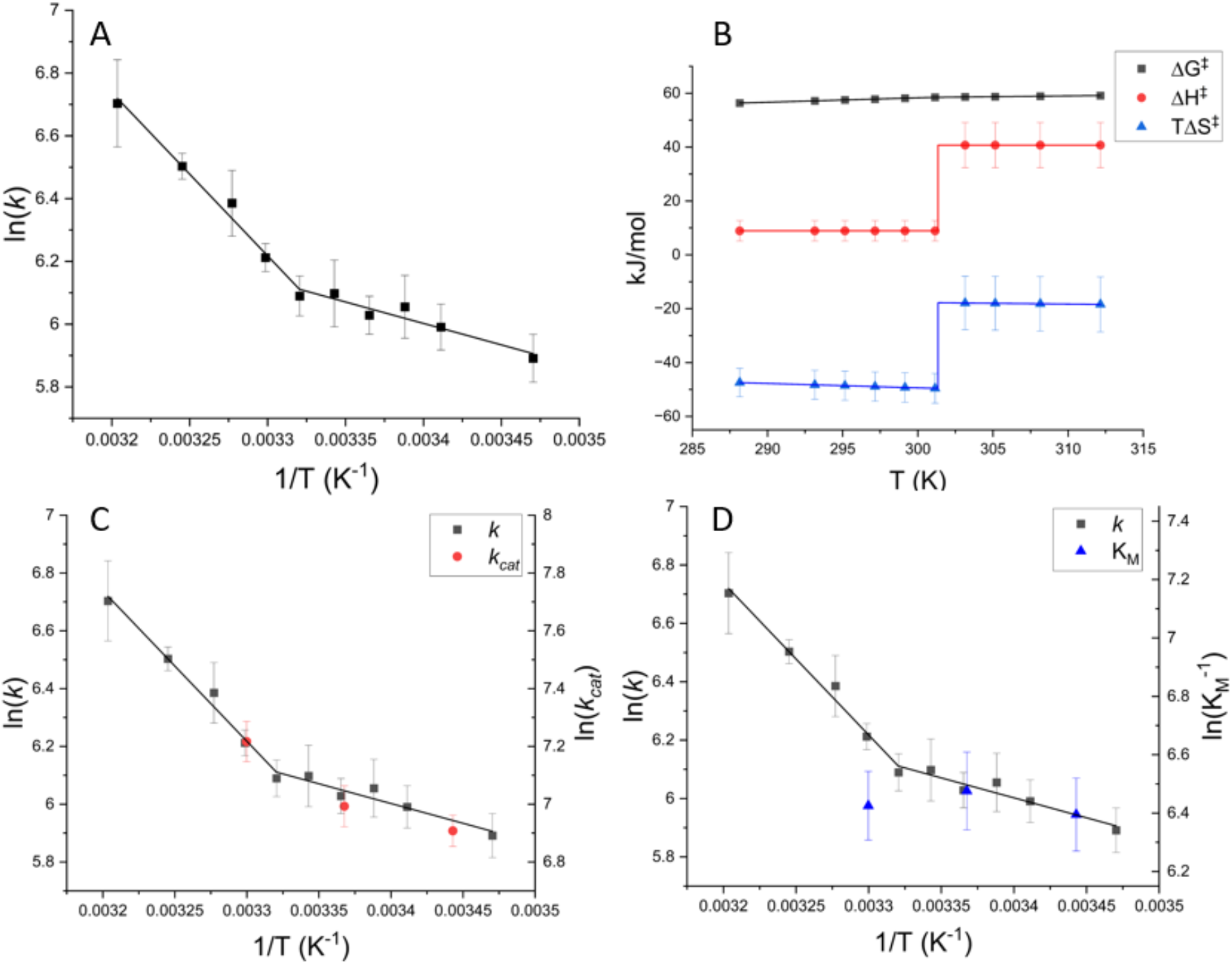
Concave Arrhenius Behavior of Thermolysin with the FAFLA Substrate. A. Thermolysin displays concave break point shaping with the FAFLA substrate, with a break point temperature of 28 °C. B. An analysis of the underlying activation energetic parameters reveals a concurrent 34 ± 8 kJ/mol increase in enthalpy (ΔH^‡^) and entropy (TΔS^‡^) of activation, which stabilizes the free energy of activation (ΔG^‡^). C. Superimposing Arrhenius-transformed *k*_*cat*_ values onto the Arrhenius plot of rate as a function of temperature shows *k*_*cat*_ changing in a concave manner, consistent with the *k*_*cat*_ being the origin of the break point. D. Superimposing Van’t Hoff-transformed K_M_ values onto the Arrhenius plot of rate as a function of temperature shows no significant changes in K_M_. The K_M_ is not sufficiently sensitive to temperature to contribute to the concave break point behavior. Error bars represent 95% confidence intervals.

### Rate-Limiting Step Determination using Kinetic Solvent Viscosity and Isotope Effects

Viscosity values for 0-30% (w/w) sucrose solutions were collected as previously described and used to evaluate KSVEs on thermolysin with the FAFLA substrate^1^. KSVEs can be examined by fitting inverse relative *k*_*cat*_ ((*k*_*cat*_)_o_/(*k*_*cat*_)_η_) as a function of relative viscosity (η_rel_), with the slope of the linear fit ranging from 0 (not sensitive to viscosity, chemical step limited) to 1 (entirely sensitive to viscosity, physical step limited)(Figure S4)^31^. Thermolysin exhibits a slope of 0.91 ± 0.10, indicating that *k*_*cat*_ is almost entirely viscosity-dependent and that chemical steps contribute minimally to *k*_*cat*_ (Figure 3A).

**Figure 3.**
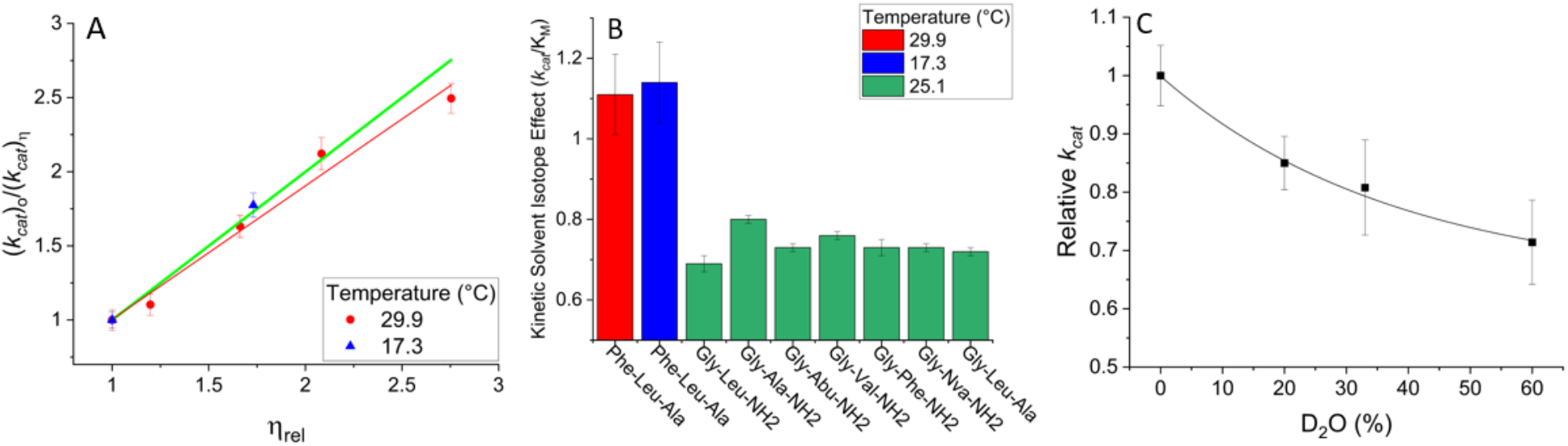
Kinetic Solvent Viscosity and Isotope Effects on Thermolysin with the FAFLA Substrate. A. Measurements of *k*_*cat*_ performed as a function of viscosity induced by additions of sucrose in the kinetic assay showed an almost complete dependence of *k*_*cat*_ on viscosity above (29.9 °C, red) and below (17.3 °C, blue) the break point. This indicated that a physical step, such as product release, is fully rate-limiting for FAFLA turnover. The green line represents the slope for a theoretical fully diffusion-sensitive enzyme^31^. B. Kinetic solvent isotope effects of D_2_O on the *k*_*cat*_/K_M_ of various N-[3-(2-furyl)acryloyl] substrates demonstrated a different isotope effect between FAFLA (red,blue) and previously-characterized, chemical step dependent substrates (green)^19,32^. C. Proton inventory of thermolysin at 29.9 °C in the presence of 0-60% D_2_O showed a modest k_cat_ decrease with a curved shape, which is consistent with viscosity effects suggesting the chemical steps are not rate limiting^33^. Error bars represent 95% confidence intervals.

KSIEs measured in 98% D_2_O on the *k*_*cat*_/K_M_ of FAFLA above (29.9 °C) and below (17.3 °C) the break point show modest, normal KSIEs of 1.11 ± 0.10 and 1.14 ± 0.10, respectively, according to Eq. 1 (Figure 3B). Previously-characterized substrates, namely FAGLA and N-[3-(2-furyl)acryloyl]-Gly-Leu-Ala that exhibit convex behavior as a thermolysin substrate, exhibit larger, inverse KSIEs around 0.70-0.80 (Figure 3B)^19,32^.

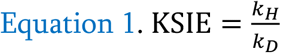

Measurements of *k*_*cat*_ (Figure 3C) and K_M_ (Figure S5) at 29.9 °C in the presence of 0-60% D_2_O demonstrated a decrease in both parameters. The shape of the proton inventory of *k*_*cat*_ appears to be exponential; suggesting a rate limiting physical step consistent with KSVEs^33^, although the proton inventory also fits to a linear model (Figure S6). Extrapolation to 100% D_2_O yields a 1.50-fold decrease in *k*_*cat*_ for the exponential fit and a 2.00-fold decrease for the linear fit. The K_M_ decrease as a function of D_2_O follows an exponential fit, which predicts a 1.45-fold decrease in K_M_ at 100% D_2_O. Combining these extrapolation values predicts *k*_*cat*_/K_M_ KSIEs at 100% D_2_O of 1.03 (*k*_*cat*_ exponential fit) and 1.38 (*k*_*cat*_ linear fit). The measured KSIE on *k*_*cat*_/K_M_ (1.11 ± 0.10) is consistent with the exponential prediction, indicating that the proton inventory follows an exponential form.

Since D_2_O is approximately 22% more viscous than H_2_O (η_rel_ of 1.22, equivalent of 11% sucrose) at 29.9 °C^34^ and the *k*_*cat*_ of thermolysin has been shown to exhibit a 91% dependence on increased viscosity with the FAFLA substrate, the observed 1.50 KSIE at 100% D_2_O can be deconvoluted to remove the effects of viscosity. This deconvolution yields a KSIE on *k*_*cat*_ of 1.25 attributed solely to the presence of deuterium (Figure S7).

### Characterization of Hinge Bending Motions using Stopped Flow Fluorescence

Since KSVEs and KSIEs suggested that a rate limiting physical step was causing the concave Arrhenius behavior for FAFLA turnover, the hinge bending motions of thermolysin were investigated for any biphasic temperature dependence. The binding of the inhibitor phosphoramidon follows the kinetic scheme in Figure S8^35–37^, where the unimolecular open-to-closed conformational change causes desolvation of the active site and the fluorescence intensity of the enzyme to increase by approximately 30%. The K_d_ for the closed enzyme-inhibitor complex is modeled by a combination of all four rate constants in Figure S8 and is present as Eq. 2, where K_sc_ approximates the equilibrium constant for the early bimolecular substrate capture step^35,36^. Fluorescence titrations using phosphoramidon and thermolysin were useful for approximating K_d_, however resulted in unacceptably large confidence intervals since K_d_<<[E_t_] (Figures S9 and S10)^35–38^. Talopeptin, which differs from phosphoramidon by one stereocenter on the deoxymannose ring, has been previously shown to exhibit a linear K_d_ (K_i_) as a function of temperature^38^: consistent with the behavior of K_d_ values for another competitive inhibitor^17^ and K_M_ values for most substrates examined^1,18^. It is less certain whether the hinge bending motions that compose the K_d_ value show break point behavior, as a linear K_d_ with temperature could mean *k*_*on*_ and *k*_*off*_ are either both linear or both show the same form of biphasic behavior.

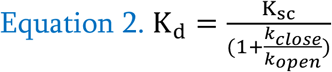

The observed rate constant for phosphoramidon binding obeys saturation kinetics (Eq. 3) due to the interplay of a fast bimolecular association and a slow unimolecular conformational change step (Figure S11)^35,36^. The existence of this unimolecular open-to-closed step was previously known for related inhibitors such as talopeptin^35^ and N-phosphoryl-Leu-Trp (PLT)^36^, but has not been confirmed for phosphoramidon^37^. It was determined that the values of *k*_*close*_ and K_sc_ were 0.48 ± 0.04 s^−1^ and 0.28 ± 0.05 mM, respectively. The closed-to-open motion, *k*_*open*_ (3.1 ×10^−5^ s^−1^), being analogous to product release, was calculated according to Eq. 2 using an estimated K_d_ of 18 nM.

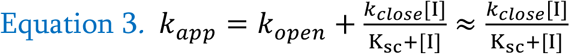

The temperature dependence of closing motions represented by 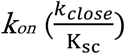 was found to be entirely linear across the 15.8-38.2 °C temperature range (Figure S12). The K_d_ is expected to be linear as well^38^, therefore the opening motions analogous to product release should also be linear with temperature. Since these conformational change steps are monophasic with temperature, they are unlikely to be rate limiting for FAFLA turnover

### Ionic Strength and Macromolecular Crowding Effects

Increasing the total ionic strength in the assay from 0.1 M (buffer alone) to 1.1 M (buffer with 1.0 M NaCl) caused the *k*_*cat*_ to increase, whereas the K_M_ for FAFLA did not change significantly (Figure 4A)(Figure S13). The ΔH°_c_ and TΔS°_c_ decreased from 34 ± 8 kJ/mol to 18 ± 6 kJ/mol between 0.1-2.5 M ionic strength, reflecting an attenuated energetic barrier and greater linear character across the break point (Figure 4B)(Figure S14). This is consistent with results obtained using the succinylcasein substrate despite the different break point shaping^1^.

**Figure 4.**
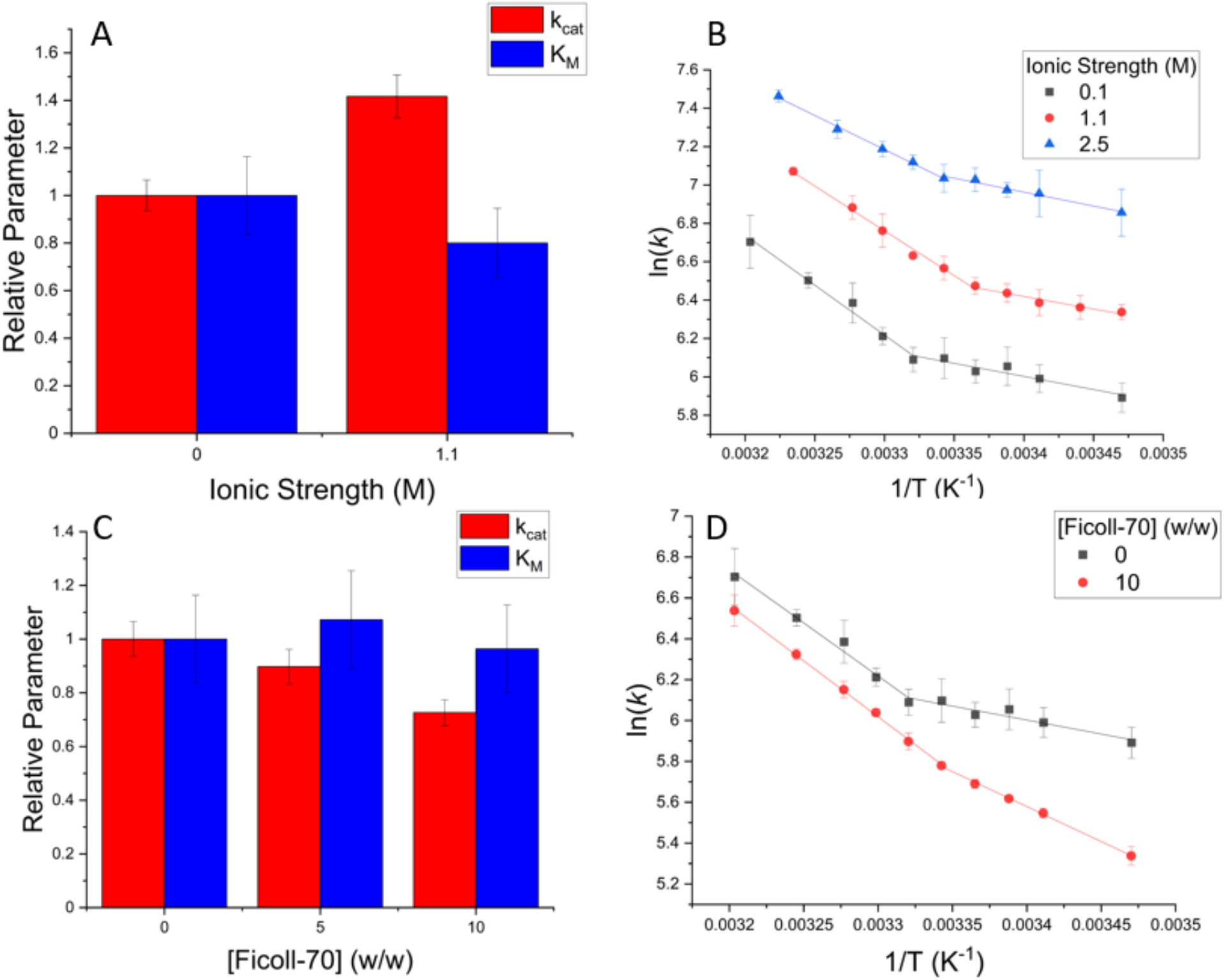
Ionic Strength and Macromolecular Crowding Effects. A. Increased concentrations of ionic strength increased the *k*_*cat*_ and did not significantly affect the K_M_ for FAFLA. B. The ΔH°_c_ and TΔS°_c_ decreased from 34 ± 8 kJ/mol to 18 ± 6 kJ/mol between 0.1-2.5 M ionic strength. C. Increased concentrations of Ficoll-70 decreased *k*_*cat*_ and did not significantly affect the K_M_ for FAFLA. D. The ΔH°_c_ and TΔS°_c_ decreased from 34 ± 8 kJ/mol to 18 ± 3 kJ/mol between 0-10% Ficoll-70. Similar to ionic strength effects, this reflects a diminished energetic severity of the break point and greater linear character of the full Arrhenius plot. Error bars represent 95% confidence intervals.

Macromolecular crowding induced by additions of the 70 kDa sucrose polymer Ficoll-70 caused the *k*_*cat*_ to decrease and the K_M_ for FAFLA to not change between 0-10% (w/w) (Figure 4C)(Figure S15). Similar to ionic strength and previous results with the succinylcasein substrate, the ΔH°_c_ and TΔS°_c_ decreased from 34 ± 8 kJ/mol to 18 ± 3 kJ/mol between 0-10% Ficoll-70, again reflecting diminished energetic severity of the break point (Figure 4D)(Figure S16)^1^.

## DISCUSSION

Thermolysin typically exhibits convex Arrhenius behavior with ΔH°_c_ and TΔS°_c_ values of approximately −30 kJ/mol^1,2,17–19^, reflecting a decrease in ΔH^‡^ and TΔS^‡^ with increasing temperature. The FAFLA substrate is novel in that thermolysin demonstrates an approximate 30 kJ/mol increase in ΔH^‡^ and TΔS^‡^ at the break point temperature near 26 °C, producing a concave Arrhenius plot. It was hypothesized that a different step in the catalytic pathway of thermolysin was rate limiting with the FAFLA substrate: leading to different break point shaping.

Previously, it was suggested that the atypical Arrhenius behavior of thermolysin could be explained by chemistry being limiting at low temperature and a physical step being limiting at high temperature, despite the authors observing no change in KSIEs across temperature^19^. Here, however, FAFLA turnover is fully limited by a physical step both above and below the break point (Figures 3A and B): the identical KSVEs and KSIEs across temperature argue against a model in which the rate limiting step switches across the break point. Variation in a single step of the pathway likely underlies the observed biphasic temperature dependence.

Figure 5A summarizes the 7 major steps of the catalytic pathway of thermolysin^19,39–42^. Step 1 is a bimolecular association of enzyme and substrate and does not contribute any rate constants to *k*_*cat*_, and therefore cannot be the origin of concavity (Figure S17)^31^. The hinge bending motions of steps 2 and 5 were found to show monophasic behavior with temperature and can be excluded assuming that hinge bending motions with phosphoramidon mirror those with FAFLA^35^. The near-complete dependence of *k*_*cat*_ on viscosity, modest KSIE on *k*_*cat*_ of ~1.25, different KSIE on the *k*_*cat*_/K_M_ and KSVEs on *k*_*cat*_ when compared to chemical step limited substrates^19,31,32,39^, and curved shape of the proton inventory indicated that the chemical steps 3 and 4 were not rate limiting with the FAFLA substrate^33^. Taken together, the data suggests that either the release of Leu-Ala (step 6) or N-[3-(2-furyl)acryloyl]-Phe (FA-Phe)(step 7) is rate limiting and the origin of concavity.

**Figure 5.**
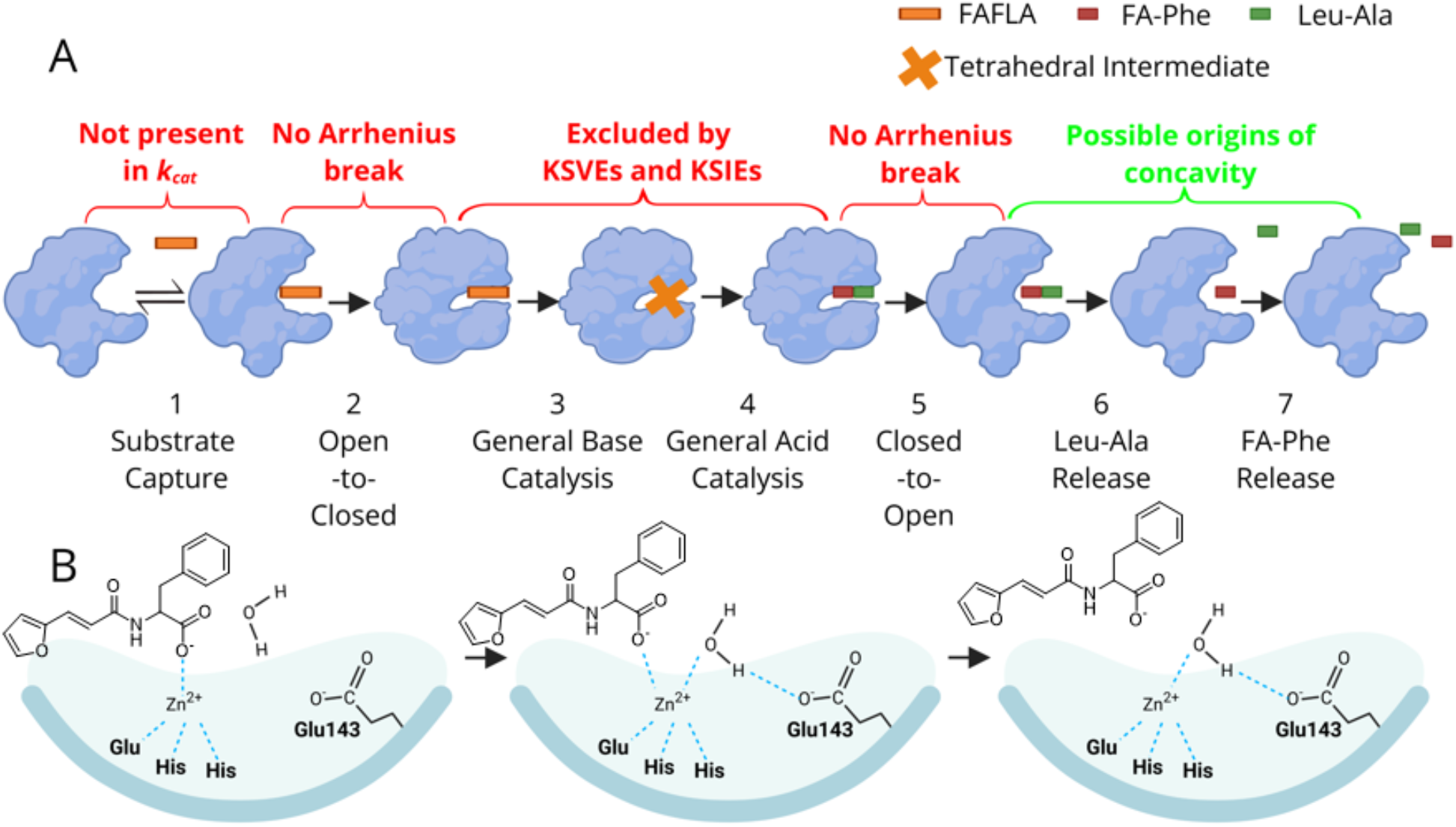
Turnover Pathway and Slow Release of FA-Phe of Thermolysin. A. The catalytic cycle of thermolysin is shown as a 7 step process^19,40–42^. Steps 1 and 2 describe substrate binding analogous to the two-step process of inhibitor binding in Figure S8. Steps 3 and 4 describe the chemical steps: nucleophilic attack followed by normally rate limiting decomposition of a tetrahedral intermediate into products^19,32^. Steps 5-7 describe product release, which is composed of a closed-to-open conformational change and sequential release of products^19,40^. The text above each step shows either which experiment excluded that step from consideration (red) or possible steps responsible for concavity (green). B. Release of FA-Phe (step 7) is likely the slower of the two product releases since FA-Phe is tethered to the enzyme through coordination to the catalytic zinc ion. FA-Phe is released when a nearby water molecule coordinates to zinc, forming a pentahedral-coordinated zinc, which then decomposes into a ground-state tetrahedral zinc with release of product^40^. Figure created using BioRender and ChemDraw 2.22.0 software.

A product release mechanism for thermolysin proposed by Hangauer *et al*. states that the N-terminal product (Leu-Ala) is released quickly due to unfavorable steric and electrostatic interactions upon amide bond cleavage, while the C-terminal product (FA-Phe) is coordinated to the catalytic Zn^2+^ and releases slowly (Figure 5B)^40^. FA-Phe release was proposed to require coordination of a nearby water molecule to Zn^2+^ to form a pentahedral-coordinated intermediate which then decomposes to a tetrahedral Zn^2+^ with concurrent release of product^40^. Other studies on thermolysin amide bond synthesis suggest that the carboxyl-containing substrate binds first^42^ or tighter^40,41^ than the amine-containing substrate: consistent with slow carboxyl-containing product release. Moreover, the unique binding mode^24^ and stability of bound complexes through interactions with Asn112 of thermolysin with FLA ligands^22,23^ are due to interactions with Phe in the P1’ position, suggesting the enzyme-product complex may also be particularly stable. Release of FA-Phe is a physical step expected to be sensitive to viscosity, and the contribution of water molecules coordinating to Zn^2+^, hydrogen bonding with Glu143, and resolvating of the active site as FA-Phe is released may explain the modest yet significant KSIE^40^.

The effects of ionic strength and macromolecular crowding on the break point of thermolysin with the FAFLA substrate were more modest but mirrored the effects seen with succinylcasein^1^: higher concentrations of NaCl and Ficoll-70 reduce ΔH°_c_ and TΔS°_c_ and increase linearity. Ionic strength and macromolecular crowding effects on the *k*_*cat*_ of thermolysin may provide some mechanistic insight into differences in rate limiting step between FAFLA and succinylcasein turnover. Thermolysin was also suggested to be primarily rate limited by a physical step with the succinylcasein substrate despite convex break point shaping^1^. FAFLA and succinylcasein exhibit different *k*_*cat*_ changes in the presence of both ionic strength and macromolecular crowding, which may be indicative of either a different rate limiting physical step or a different mechanism of product release between small molecule and macromolecular substrates. Furthermore, the size of cleavage products affecting diffusion rate, potential for downstream interactions, and unknown cleavage sites (if Phe in P1’ is important for concavity) may contribute to different break point shaping with the succinylcasein substrate.

It was previously observed that macromolecular crowding exerted a more modest effect on the activation energetic parameters of the low temperature regime by 4-fold (Figure 6A), which resulted in the decreased ΔH°_c_ and TΔS°_c_ and may be interpreted to mean that the thermolysin conformation or ensemble of the low temperature regime was stabilized or favored in a crowded environment^1^. This effect was reversed for the concave Arrhenius plot, where the E_a_ of the high temperature regime was not significantly affected and the low temperature regime demonstrated an E_a_ increase (Figure 6B)^1^. A similar switch occurred for ionic strength (Table S1). This is not consistent with the model that a particular set of conformers is favored or stabilized in each environment, as identical effects would be expected in this case. It is consistent that the high E_a_ (high ΔH^‡^ and TΔS^‡^) phase is less sensitive to macromolecular crowding and the low E_a_ (low ΔH^‡^ and TΔS^‡^) phase is less sensitive to ionic strength^1^. This suggests that differences in activation parameters across the break point temperature may be driving Arrhenius linearization rather than stabilization of structural features.

**Figure 6.**
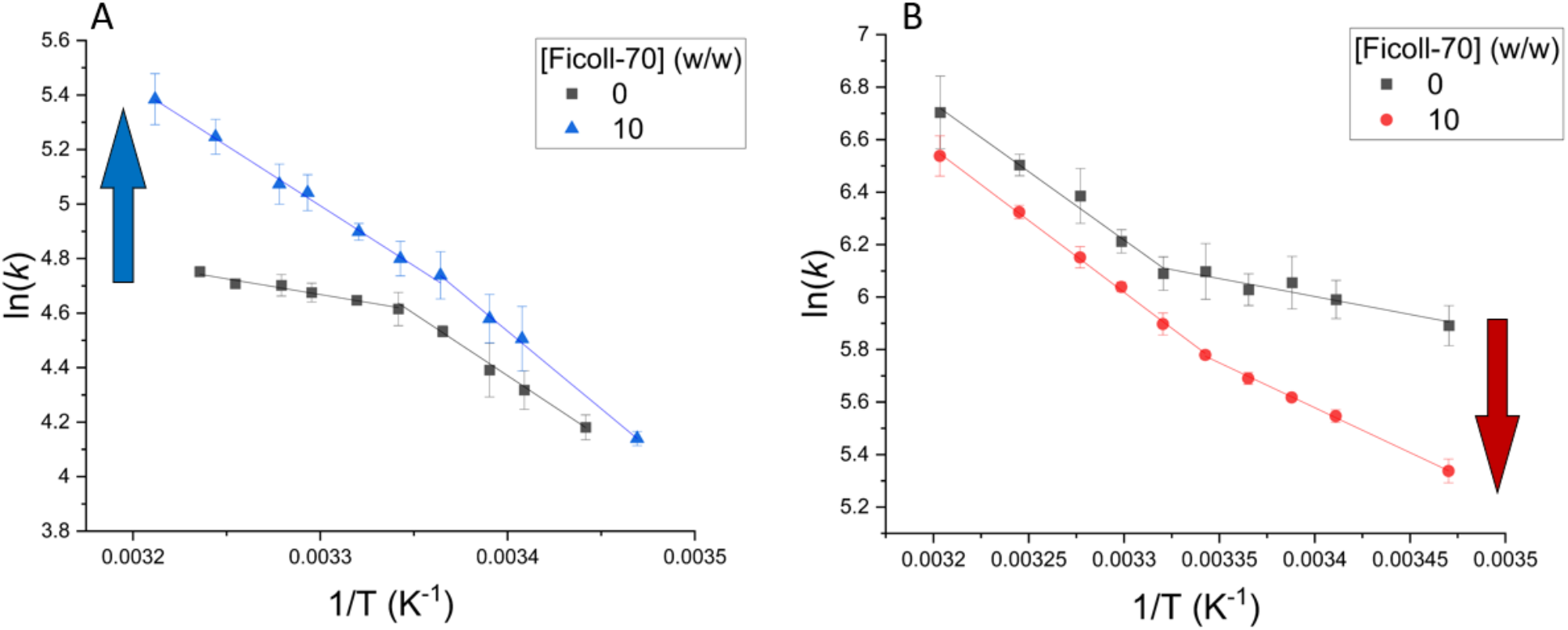
Macromolecular Crowding Affects Different Temperature Regimes of the Arrhenius Plot Between Succinylcasein and FAFLA Substrates. A. The high temperature regime was linearized relative to the low temperature regime with the succinylcasein substrate with added macromolecular crowding^1^. B. This effect was reversed with the FAFLA substrate, where the low temperature regime was linearized relative to the high temperature regime. A similar change occurred in elevated ionic strength. Error bars represent 95% confidence intervals.

While rare in kinetic temperature dependences, concave behavior is common when examining break point phenomena through structural methods. For example, *Geobacillus stearothermophilus* alcohol dehydrogenase (htADH, convex temperature dependence of chemical steps)^3,43^, ribonuclease A (RNase A)^44^, and myoglobin^45,46^ all indicate an abrupt increase in structural dynamics at temperatures above their respective break points consistent with the increase in entropy and enthalpy with a concave kinetic break point. Given that the concave break point of thermolysin is linked to a physical process, with ΔH°_c_ and TΔS°_c_ possibility reflecting an enzyme conformational change at the break point^30^ consistent with those observed in the above three proteins, this shape may directly report on the dynamic state of the enzyme, rather than reflecting how dynamics modulate chemistry. Overall, the relationship between physical and chemical rate limitation and break point shape remains weakly characterized because few enzymes have been examined (Table 1)^1–14,19,25–27,47–56^; however, the few examples of concave Arrhenius behavior thus far have only been observed with physical step limitation of *k*_*cat*_: indicating some correlation.

**Table 1.**
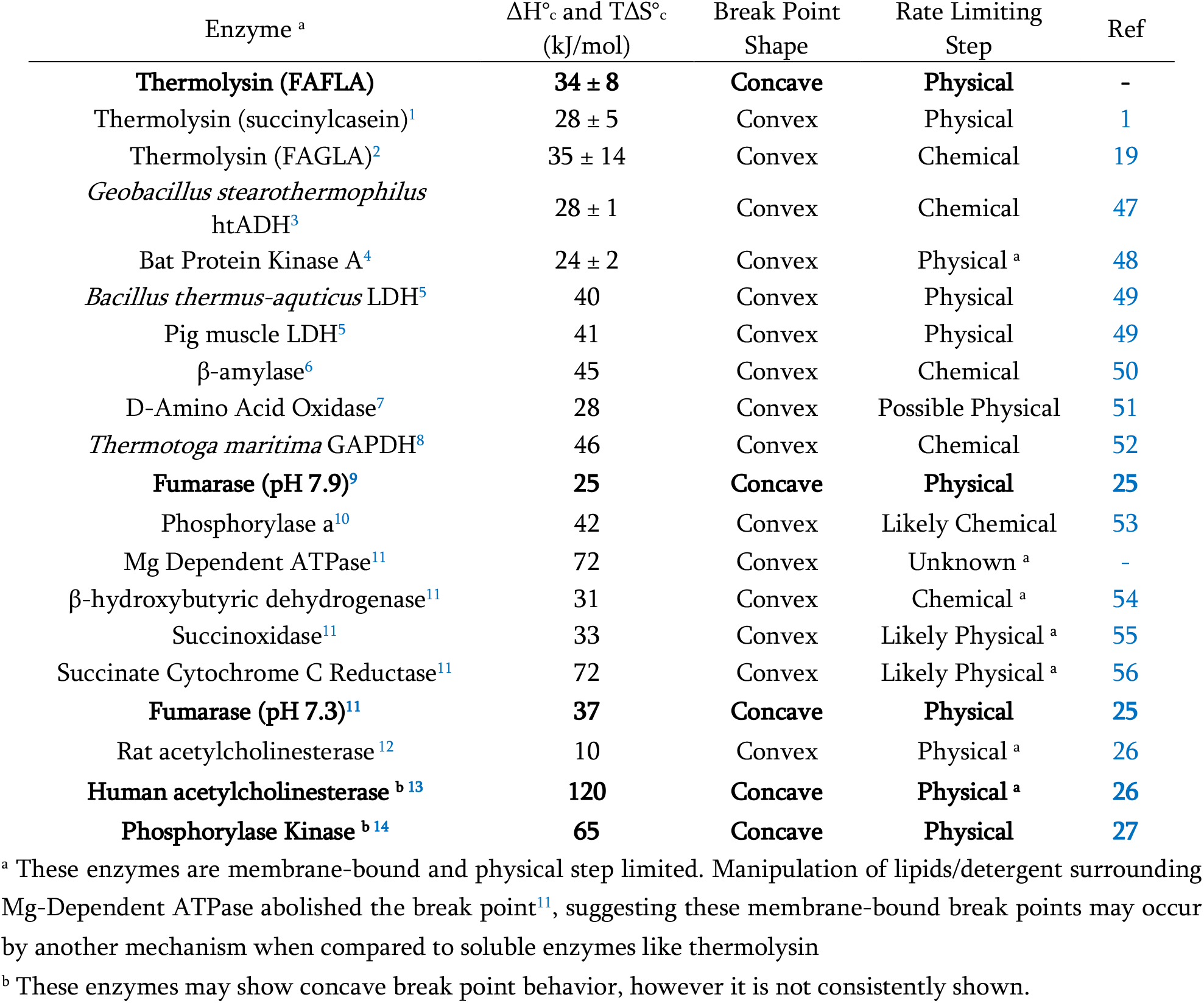
Previously-Characterized Enzymes with Break Point Behavior and Rate Limiting Step.

## MATERIALS AND METHODS

### Standard Kinetic Assay

Steady-state kinetics experiments were performed on an HP-8453 UV-Vis spectrophotometer with a temperature-controlled cuvette holder connected to an external water bath. Temperature was measured using a digital thermocouple. FAFLA was synthesized by Biomatik while thermolysin was supplied as the type X powder (Sigma). Concentrations of FAFLA and thermolysin were ascertained using ε_305_ of 17.9 mM^−1^cm^−1^ (Figure S18) and ε_280_ of 61.1 mM^−1^cm^−1^, respectively.

Enzyme and substrate samples were allowed to equilibrate to temperature in the water bath for 3 min prior to mixing. When applicable, FAFLA and thermolysin were dissolved separately in buffer containing the desired amount of ionic strength, viscosity, or macromolecular crowding to promote mixing. The assay consisted of 900 µL of FAFLA dissolved in 50 mM sodium borate buffer to produce final concentrations of 0-3 mM and 100 µL of thermolysin dissolved in 50 mM 2-(N-morpholino)ethanesulfonic acid (MES) (pH 7.50) with a final concentration of 1-2 nM. Experiments characterizing Arrhenius behavior and ionic strength/macromolecular crowding effects used pH 7.20 borate buffer whereas KSVEs with added sucrose were performed at pH 8.00. This change in pH did not affect the break point energetics (Figure S19). Cleavage of FAFLA was measured continuously at 350 nm with baseline correction at 700 nm for 90 s with a spectrum recorded every 1 s. Rates were obtained from initial rate fits over the time course and converted to concentrations using Δε_350_ of −0.104 mM^−1^cm^−1^ (Figures S20-S21). This process was repeated for each concentration of substrate or temperature in triplicate. Steady-state kinetic parameters were calculated in Origin v.2024 software using a non-linear-least-squares fit to a Michaelis-Menten model (Eq. 4). Arrhenius parameters were calculated as previously described^1^, with an initial piecewise fitting (PWL2, Eq 5-7) to determine the break point temperature followed by linear fits to each regime to attain E_a_ values (Eq. 8). Kinetic temperature dependences of thermolysin were performed at a fixed FAFLA concentration of 0.70 mM and ΔH°_c_ and TΔS°_c_ values, which refer to the difference in enthalpy of activation (ΔH^‡^) and entropy of activation (TΔS^‡^)^30^ across the break point, were obtained using Eq. 9. The fitting model used assumes that the two phases of the Arrhenius plot meet at the same temperature and confers perfect entropy-enthalpy compensation, which is why ΔH°_c_ and TΔS°_c_ are equivalent at the break point temperature.

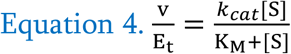

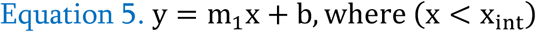

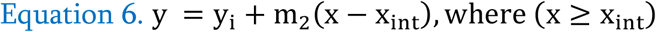

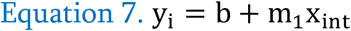

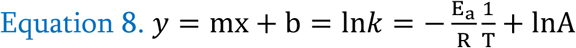

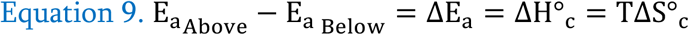

### Viscometry for Kinetic Solvent Viscosity Effects

Viscosity values of 0-30% (w/w) sucrose solutions were obtained as previously described on a temperature-controlled Anton Paar AMVn falling ball viscometer^1^.

### Deuteration of Buffer and Kinetic Solvent Isotope Effects

An aliquot of 50 mL of 50 mM sodium borate buffer (pH 7.50) was frozen and lyophilized until the pressure reading was <25 mTorr, indicating complete removal of H_2_O. The dry buffer was then quickly redissolved in 50 mL of D_2_O, sealed with parafilm, and lightly shaken until fully dissolved. The pD of this deuterated sodium borate buffer was found to be 8.07 according to Eq. 10 where pH_a_ is the pH electrode reading^57^. The buffer was stored in 10 mL aliquots at 4 °C for 5 d.

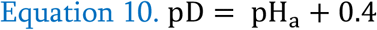

KSIEs on the *k*_*cat*_/K_M_ of thermolysin with the FAFLA substrate were determined using 980 µL of the above deuterated buffer and 10 µL of FAFLA (69 µM) and thermolysin (1 nM) each in protonated 50 mM sodium borate buffer (pH 7.50)(Figure S22) on a Cary 300 UV-Vis with Peltier temperature control. Measurements of *k*_*cat*_ and K_M_ to construct a proton inventory were performed using the standard kinetic assay using combinations of buffer with pH 7.50 and pD 8.07 (Figure S23).

### Stopped Flow Fluorescence of Phosphoramidon Binding Kinetics

Stopped flow fluorescence measurements were performed as an adaptation of a published procedure on a Hi-Tech Scientific SF-61DX2 Stopped Flow Spectrophotometer^35–38^ with instrument temperature maintained by an external water bath. Thermolysin and phosphoramidon (ε_280_ of 5.55 mM^−1^cm^−1^) were separately dissolved in 50 mM sodium borate buffer (pH 8.00) to final concentrations of 2.3 µM and 0-0.5 mM, respectively. The excitation wavelength was set to 280 nm with emission measured with a cut-off filter at 320 nm. Samples were injected using a drive pressure of 5 kgf/cm^2^ (0.5 MPa) with a dead time of 1.8 ± 0.5 ms determined using tryptophan and N-bromosuccinimide in triplicate (Figure S24). Fluorescence was recorded over the time course which ranged from 30-120 seconds (Figure S25). The time courses were fit using a single exponential in Kinetic Studio software to obtain the value of *k*_*app*_. Time courses were performed in triplicate for each concentration of inhibitor or temperature for each experiment.

The behavior of phosphoramidon binding to thermolysin follows Eq. 3^35,36^, which is a modified hyperbolic equation offset from the origin by the back-reaction constant *k*_*open*_. Since *k*_*open*_ *<< k*_*close*_ by 15,000-fold, it was difficult to obtain accurate values of *k*_*open*_ from the fit, therefore the equation was fit to a standard hyperbolic Michaelis-Menten model with the asymptote being *k*_*close*_. Values of *k*_*open*_ were estimated according to Eq. 2, using values of *k*_*close*_ and K_sc_ from the fit and estimated value of K_d_ at the appropriate temperature from the fluorescence titration.

Starting at inhibitor concentrations 10-fold lower than K_sc_, the value of 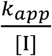 approximates the value of 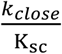. The Arrhenius plot of 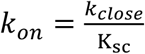 were determined at a fixed phosphoramidon concentration of 17 µM.

## ABBREVIATIONS

AchE: acetylcholinesterase
E_a_: activation energy
FAFLA: N-[3-(2-furyl)acryloyl]-Phe-Leu-Ala
FAGLA: N-[3-(2-furyl)acryloyl]-Gly-Leu-Amide
FA-Phe: N-[3-(2-furyl)acryloyl]-Phe
FLA: Phe-Leu-Ala
ΔG^‡^: free energy of activation
ΔH°_c_: enthalpy difference across the break point
ΔH^‡^: enthalpy of activation
htADH: *Geobacillus stearothermophilus* alcohol dehydrogenase
KSIE: Kinetic Solvent Isotope Effect
KSVE: Kinetic Solvent Viscosity Effect
MES: 2-(N-morpholino)ethanesulfonic acid
PhK: phosphorylase kinase
PLT: N-phosphoryl-Leu-Trp
RNase A: ribonuclease A
TΔS^‡^: entropy of activation
succinylcasein: succinylated bovine β-casein
TΔS°_c_: entropy difference across the break point
ZFLA: N-benzyloxycarbonyl-Phe-Leu-Ala
ZF^P^LA: N-benzyloxycarbonyl-Phe-phosphoryl-Leu-Ala
ZG^P^LA: N-benzyloxycarbonyl-Gly-phosphoryl-Leu-Ala

## SUPPLEMENTARY MATERIAL

A separate Supplementary Information File contains raw kinetic data, values used in the construction of Figures, and supplementary data to support claims made in the article.

## ACKNOWLEDGMENTS

We would like to thank the Delaware Biotechnology Institute for the use of their Agilent Eclipse Fluorometer and Cary 300 UV-Vis, Dr. Norman Wagner and Sean Farrington for the use of their Anton Paar viscometer, and Dr. Joseph M. Fox and Will Trout for the use of their Hi-Tech Scientific SF-61DX2 Stopped Flow Spectrophotometer. Origin v.2024 software was used to generate figures and BioRender software was used in Figure 5. Funding for this article was provided by the Department of Chemistry and Biochemistry at the University of Delaware.

## DATA AVAILABILITY

The data that supports the findings of this study are available in the supplementary material of this article (Tables S2-S34). The data that support the findings of this study are available from the corresponding author upon reasonable request.

