## Supplementary Materials for "Rate limiting release of product underlies concave Arrhenius break point of thermolysin with a Phe-Leu-Ala substrate"

Supporting Information

**Table S1**

| | Succinylcasein (Convex) | | FAFLA (Concave) | |
|---|---|---|---|---|
| Ionic Strength (M) | $E_a$ Above Break (kJ/mol) | $E_a$ Below Break (kJ/mol) | $E_a$ Above Break (kJ/mol) | $E_a$ Below Break (kJ/mol) |
| 0.1 | 10.0 ± 2.5 | 38.2 ± 4.2 | 44.9 ± 6.7 | 10.8 ± 4.2 |
| 0.35 | 7.7 ± 1.2 | 28.3 ± 2.5 | - | - |
| 0.6 | 7.2 ± 2.7 | 23.3 ± 6.7 | - | - |
| 0.85 | 5.5 ± 1.8 | 18.3 ± 1.7 | - | - |
| 1.1 | 2.8 ± 2.1 | 10.8 ± 0.8 | 39.1 ± 5.0 | 10.8 ± 3.3 |
| 2.5 | - | - | 29.9 ± 3.3 | 12.5 ± 5.0 |

**Table S1.** Activation energy ($E_a$) changes of thermolysin across the break point in elevated ionic strength show that $E_a$ below the break is affected more by ionic strength with succinylcasein and $E_a$ above the break is affected more by ionic strength with the FAFLA substrate. This mirrored the effects observed with elevated macromolecular crowding.

**Table S2**

| 1/T ($K^{-1}$) | Rate ($s^{-1}$) | | |
|---|---|---|---|
| 0.003470 | 378.5 | 333.9 | 373.4 |
| 0.003411 | 370.3 | 410.4 | 418.5 |
| 0.003388 | 455.5 | 383.8 | 439.7 |
| 0.003365 | 423.3 | 390.0 | 432.0 |
| 0.003343 | 451.3 | 400.2 | 483.0 |
| 0.003321 | 465.0 | 443.4 | 415.5 |
| 0.003299 | 516.9 | 501.5 | 478.0 |
| 0.003277 | 654.8 | 549.7 | 574.9 |
| 0.003245 | 695.0 | 651.9 | 654.7 |
| 0.003204 | 930.5 | 758.0 | 757.5 |

**Table S2.** Raw data collected at 0.70 mM FAFLA at pH 7.20 in the absence of elevated ionic strength or macromolecular crowding.

**Table S3**

| 1/T ($K^{-1}$) | Rate ($s^{-1}$) | | |
|---|---|---|---|
| 0.003470 | 305.1 | 320.6 | 290.2 |
| 0.003411 | 348.5 | 341.4 | 344.4 |
| 0.003388 | 367.6 | 337.5 | 363.6 |
| 0.003365 | 367.8 | 373.2 | 384.8 |
| 0.003343 | 389.4 | 411.8 | 392.7 |
| 0.003320 | 438.2 | 453.6 | 451.6 |
| 0.003298 | 481.0 | 512.2 | 481.5 |
| 0.003277 | 565.6 | 577.4 | 583.4 |
| 0.003245 | 666.6 | 691.4 | 733.2 |

**Table S3.** Raw data collected at 0.70 mM FAFLA at pH 8.00 in the absence of elevated ionic strength or macromolecular crowding.

**Table S4**

| 1/T ($K^{-1}$) | Rate ($s^{-1}$) | | |
|---|---|---|---|
| 0.003470 | 555.0 | 587.9 | 553.4 |
| 0.003440 | 562.1 | 615.6 | 560.4 |
| 0.003411 | 567.9 | 634.5 | 578.5 |
| 0.003388 | 608.1 | 654.5 | 610.9 |
| 0.003365 | 635.4 | 677.3 | 632.8 |
| 0.003343 | 705.9 | 751.3 | 675.6 |
| 0.003320 | 753.2 | 761.1 | 763.1 |
| 0.003298 | 800.3 | 859.2 | 932.2 |
| 0.003277 | 921.6 | 1026.6 | 976.6 |
| 0.003234 | 1154.3 | 1185.6 | 1192.8 |

**Table S4.** Raw data collected at 0.70 mM FAFLA at pH 7.20 in the presence of 1.1 M ionic strength.

**Table S5**

| 1/T ($K^{-1}$) | Rate ($s^{-1}$) | | |
|---|---|---|---|
| 0.003470 | 831.3 | 1001.1 | 1016.4 |
| 0.003411 | 1079.4 | 1143.0 | 924.7 |
| 0.003388 | 1067.2 | 1034.2 | 1104.7 |
| 0.003365 | 1086.7 | 1098.4 | 1197.8 |
| 0.003343 | 1210.7 | 1065.9 | 1128.3 |
| 0.003320 | 1277.2 | 1195.7 | 1232.9 |
| 0.003298 | 1376.7 | 1304.3 | 1286.3 |
| 0.003266 | 1413.7 | 1450.4 | 1533.2 |
| 0.003224 | 1765.8 | 1684.3 | 1776.6 |

**Table S5.** Raw data collected at 0.70 mM FAFLA at pH 7.20 in the presence of 2.5 M ionic strength.

**Table S6**

| 1/T ($K^{-1}$) | Rate ($s^{-1}$) | | |
|---|---|---|---|
| 0.003470 | 215.3 | 209.9 | 199.0 |
| 0.003411 | 262.7 | 252.8 | 253.8 |
| 0.003388 | 272.7 | 274.6 | 278.7 |
| 0.003365 | 298.5 | 300.2 | 289.4 |
| 0.003343 | 327.8 | 324.6 | 318.9 |
| 0.003320 | 351.2 | 378.2 | 363.0 |
| 0.003298 | 423.6 | 423.4 | 411.1 |
| 0.003277 | 453.6 | 467.5 | 487.1 |
| 0.003245 | 561.3 | 568.3 | 543.7 |
| 0.003203 | 652.0 | 677.5 | 742.9 |

**Table S6.** Raw data collected at 0.70 mM FAFLA at pH 7.20 in the presence of 10% Ficoll-70.

**Table S7**

| FAFLA (mM) | Rate ($s^{-1}$) | | |
|---|---|---|---|
| 0.00 | 3.6 | 2.0 | -5.6 |
| 0.11 | 141.8 | 141.6 | 138.3 |
| 0.22 | 244.8 | 252.4 | 251.4 |
| 0.33 | 345.5 | 313.7 | 334.6 |
| 0.43 | 428.6 | 443.0 | 443.2 |
| 0.54 | 541.3 | 460.0 | 485.5 |
| 0.81 | 626.2 | 607.8 | 610.3 |
| 1.08 | 590.6 | 684.1 | 626.7 |
| 2.17 | 765.4 | 738.3 | 740.7 |

**Table S7.** Raw Michaelis-Menten data collected at 29.9 °C at pH 7.20 in the absence of elevated ionic strength, viscosity, or macromolecular crowding.

**Table S8**

| FAFLA (mM) | Rate ($s^{-1}$) | | |
|---|---|---|---|
| 0.00 | -53.3 | -0.1 | 53.3 |
| 0.13 | 378.8 | 383.3 | 271.0 |
| 0.27 | 417.7 | 524.0 | 585.2 |
| 0.40 | 562.2 | 617.8 | 620.3 |
| 0.53 | 791.5 | 804.2 | 685.7 |
| 0.67 | 860.7 | 808.8 | 807.7 |
| 1.00 | 931.3 | 1001.6 | 919.9 |
| 1.33 | 1073.8 | 1057.2 | 970.8 |
| 2.66 | 1117.9 | 1132.8 | 1237.8 |

**Table S8.** Raw Michaelis-Menten data collected at 29.9 °C at pH 7.20 in the presence of 1.1 M ionic strength.

**Table S9**

| FAFLA (mM) | Rate ($s^{-1}$) | | |
|---|---|---|---|
| 0.00 | -2.4 | 0.1 | 2.3 |
| 0.11 | 202.5 | 154.9 | 125.8 |
| 0.21 | 236.3 | 212.5 | 242.5 |
| 0.32 | 302.1 | 291.5 | 287.7 |
| 0.43 | 326.7 | 337.0 | 330.3 |
| 0.53 | 414.5 | 411.1 | 392.4 |
| 0.80 | 504.8 | 522.0 | 475.0 |
| 1.07 | 593.4 | 596.4 | 569.1 |
| 2.13 | 715.0 | 643.7 | 633.1 |

**Table S9.** Raw Michaelis-Menten data collected at 29.9 °C at pH 7.20 in the presence of 5% Ficoll-70.

**Table S10**

| FAFLA (mM) | Rate ($s^{-1}$) | | |
|---|---|---|---|
| 0.00 | 9.7 | -0.2 | -9.4 |
| 0.12 | 132.4 | 189.0 | 159.0 |
| 0.24 | 195.0 | 195.8 | 210.0 |
| 0.37 | 296.1 | 317.9 | 284.2 |
| 0.49 | 339.4 | 328.7 | 296.1 |
| 0.61 | 383.1 | 346.4 | 386.4 |
| 0.92 | 421.3 | 460.0 | 415.7 |
| 1.22 | 509.9 | 506.1 | 512.4 |
| 2.45 | 564.1 | 604.1 | 547.7 |

**Table S10.** Raw Michaelis-Menten data collected at 29.9 °C at pH 7.20 in the presence of 10% Ficoll-70.

**Table S11**

| FAFLA (mM) | Rate ($s^{-1}$) | | |
|---|---|---|---|
| 0.00 | -29.9 | 14.1 | 15.7 |
| 0.18 | 144.6 | 190.1 | 139.7 |
| 0.36 | 246.4 | 251.1 | 268.7 |
| 0.53 | 346.4 | 352.4 | 309.4 |
| 0.71 | 403.7 | 423.7 | 393.6 |
| 0.89 | 479.1 | 478.9 | 461.4 |
| 1.34 | 592.4 | 572.1 | 650.0 |
| 1.78 | 734.2 | 714.8 | 745.9 |
| 3.56 | 949.7 | 870.1 | 985.0 |

**Table S11.** Raw Michaelis-Menten data collected at 29.9 °C at pH 8.00 in the absence of ionic strength, viscosity, or macromolecular crowding.

**Table S12**

| FAFLA (mM) | Rate ($s^{-1}$) | | |
|---|---|---|---|
| 0.00 | -9.2 | -0.8 | 10.0 |
| 0.19 | 147.5 | 118.8 | 138.2 |
| 0.37 | 192.7 | 213.0 | 186.1 |
| 0.56 | 284.9 | 283.8 | 242.9 |
| 0.74 | 364.7 | 304.0 | 372.6 |
| 0.93 | 408.3 | 438.6 | 437.6 |
| 1.39 | 543.7 | 555.6 | 555.2 |
| 1.85 | 622.4 | 604.4 | 561.6 |
| 3.71 | 757.1 | 750.5 | 786.4 |

**Table S12.** Raw Michaelis-Menten data collected at 23.8 °C at pH 8.00 in the absence of ionic strength, viscosity, or macromolecular crowding.

**Table S13**

| FAFLA (mM) | Rate ($s^{-1}$) | | |
|---|---|---|---|
| 0.00 | -8.0 | -12.4 | 20.4 |
| 0.18 | 121.1 | 114.2 | 91.8 |
| 0.36 | 175.6 | 178.5 | 180.5 |
| 0.54 | 229.8 | 241.9 | 244.4 |
| 0.72 | 280.3 | 299.6 | 276.2 |
| 0.90 | 348.3 | 368.6 | 342.5 |
| 1.34 | 458.8 | 440.1 | 443.0 |
| 1.79 | 530.1 | 529.7 | 541.8 |
| 3.58 | 711.2 | 666.7 | 653.4 |

**Table S13.** Raw Michaelis-Menten data collected at 17.3 °C at pH 8.00 in the absence of ionic strength, viscosity, or macromolecular crowding.

**Table S14**

| FAFLA (mM) | Rate ($s^{-1}$) | | |
|---|---|---|---|
| 0.00 | -0.5 | -17.4 | 17.9 |
| 0.17 | 156.3 | 167.1 | 153.2 |
| 0.34 | 282.0 | 245.5 | 269.8 |
| 0.51 | 354.3 | 387.4 | 340.3 |
| 0.67 | 459.3 | 421.4 | 429.5 |
| 0.84 | 498.5 | 472.6 | 514.0 |
| 1.26 | 605.5 | 591.6 | 570.0 |
| 1.68 | 773.8 | 699.6 | 758.7 |
| 3.37 | 844.2 | 944.1 | 908.2 |

**Table S14.** Raw Michaelis-Menten data collected at 29.9 °C at pH 8.00 in the presence of 10% sucrose.

**Table S15**

| FAFLA (mM) | Rate ($s^{-1}$) | | |
|---|---|---|---|
| 0.00 | -16.7 | -6.1 | 22.8 |
| 0.16 | 141.8 | 140.8 | 154.0 |
| 0.31 | 255.2 | 249.6 | 253.4 |
| 0.47 | 282.9 | 309.8 | 314.4 |
| 0.62 | 394.8 | 322.7 | 330.9 |
| 0.78 | 397.1 | 408.9 | 422.1 |
| 1.17 | 510.1 | 508.7 | 493.3 |
| 1.56 | 550.3 | 545.7 | 563.7 |
| 3.12 | 652.1 | 665.4 | 681.9 |

**Table S15.** Raw Michaelis-Menten data collected at 29.9 °C at pH 8.00 in the presence of 20% sucrose.

**Table S16**

| FAFLA (mM) | Rate ($s^{-1}$) | | |
|---|---|---|---|
| 0.00 | 3.6 | 1.9 | -5.6 |
| 0.12 | 86.2 | 95.5 | 104.9 |
| 0.24 | 146.3 | 159.2 | 173.1 |
| 0.36 | 189.3 | 165.8 | 225.8 |
| 0.48 | 251.6 | 256.4 | 246.1 |
| 0.60 | 289.2 | 284.0 | 283.0 |
| 0.90 | 329.2 | 333.0 | 342.4 |
| 1.20 | 398.7 | 385.6 | 383.0 |
| 2.39 | 490.1 | 481.2 | 500.8 |

**Table S16.** Raw Michaelis-Menten data collected at 29.9 °C at pH 8.00 in the presence of 25% sucrose.

**Table S17**

| FAFLA (mM) | Rate ($s^{-1}$) | | |
|---|---|---|---|
| 0.00 | -2.4 | -4.5 | 6.9 |
| 0.16 | 112.1 | 111.1 | 100.2 |
| 0.31 | 176.7 | 168.0 | 193.3 |
| 0.47 | 231.4 | 217.4 | 223.7 |
| 0.62 | 242.2 | 244.0 | 266.8 |
| 0.78 | 293.7 | 302.9 | 308.7 |
| 1.17 | 365.3 | 355.6 | 355.8 |
| 1.56 | 393.1 | 400.4 | 379.4 |
| 3.11 | 454.9 | 423.4 | 456.0 |

**Table S17.** Raw Michaelis-Menten data collected at 29.9 °C at pH 8.00 in the presence of 30% sucrose.

**Table S18**

| FAFLA (mM) | Rate ($s^{-1}$) | | |
|---|---|---|---|
| 0.00 | 2.1 | 18.6 | -20.7 |
| 0.17 | 109.2 | 111.8 | 121.5 |
| 0.34 | 180.1 | 153.6 | 175.7 |
| 0.50 | 230.0 | 228.2 | 215.3 |
| 0.67 | 267.7 | 258.3 | 254.5 |
| 0.84 | 296.1 | 315.9 | 309.8 |
| 1.26 | 372.3 | 369.9 | 350.7 |
| 1.68 | 385.7 | 425.7 | 386.5 |
| 3.35 | 462.9 | 442.1 | 463.4 |

**Table S18.** Raw Michaelis-Menten data collected at 17.3 °C at pH 8.00 in the presence of 20% sucrose.

**Table S19**

| FAFLA (mM) | Rate ($s^{-1}$) | | |
|---|---|---|---|
| 0.00 | -4.6 | 21.2 | -16.6 |
| 0.16 | 138.8 | 165.1 | 160.4 |
| 0.33 | 279.3 | 290.1 | 294.2 |
| 0.49 | 390.1 | 373.9 | 404.0 |
| 0.65 | 444.9 | 476.1 | 471.4 |
| 0.81 | 571.2 | 571.1 | 543.5 |
| 1.22 | 732.9 | 722.1 | 706.1 |
| 1.63 | 777.0 | 739.4 | 822.4 |
| 3.25 | 1008.9 | 928.9 | 982.3 |

**Table S19.** Raw Michaelis-Menten data collected at 29.9 °C at pH 7.50 in the presence of 0% $D_2O$.

**Table S20**

| FAFLA (mM) | Rate ($s^{-1}$) | | |
|---|---|---|---|
| 0.00 | 3.8 | -7.5 | 3.6 |
| 0.16 | 182.7 | 171.0 | 218.2 |
| 0.32 | 299.6 | 302.7 | 312.6 |
| 0.48 | 386.2 | 388.0 | 398.0 |
| 0.63 | 443.2 | 479.5 | 454.1 |
| 0.79 | 588.7 | 560.9 | 537.2 |
| 1.19 | 676.0 | 666.6 | 637.7 |
| 1.59 | 734.4 | 768.3 | 741.8 |
| 3.17 | 892.4 | 869.2 | 887.4 |

**Table S20.** Raw Michaelis-Menten data collected at 29.9 °C at pH 7.50 and pD 8.07 in the presence of 20% $D_2O$.

**Table S21**

| FAFLA (mM) | Rate ($s^{-1}$) | | |
|---|---|---|---|
| 0.00 | 7.0 | -12.2 | 5.2 |
| 0.15 | 193.4 | 168.4 | 147.0 |
| 0.30 | 292.2 | 282.4 | 304.3 |
| 0.46 | 377.8 | 409.5 | 374.1 |
| 0.61 | 382.4 | 381.2 | 460.3 |
| 0.76 | 514.3 | 515.2 | 437.0 |
| 1.14 | 665.2 | 628.9 | 634.6 |
| 1.52 | 704.4 | 717.4 | 724.7 |
| 3.05 | 826.0 | 820.5 | 829.4 |

**Table S21.** Raw Michaelis-Menten data collected at 29.9 °C at pH 7.50 and pD 8.07 in the presence of 33% $D_2O$.

**Table S22**

| FAFLA (mM) | Rate ($s^{-1}$) | | |
|---|---|---|---|
| 0.00 | -2.8 | -0.5 | 3.4 |
| 0.16 | 172.1 | 162.9 | 128.3 |
| 0.31 | 268.6 | 245.8 | 234.4 |
| 0.47 | 378.0 | 335.1 | 370.8 |
| 0.63 | 464.3 | 458.9 | 443.7 |
| 0.79 | 469.0 | 505.5 | 497.1 |
| 1.18 | 566.9 | 566.3 | 586.4 |
| 1.57 | 662.4 | 674.9 | 685.6 |
| 3.14 | 743.5 | 732.1 | 763.1 |

**Table S22.** Raw Michaelis-Menten data collected at 29.9 °C at pH 7.50 and pD 8.07 in the presence of 60% $D_2O$.

**Table S23**

| Solvent | Approximated $k_{cat}/K_M$ ($\mu M^{-1}s^{-1}$) | | | | | Average ($\mu M^{-1}s^{-1}$) |
|---|---|---|---|---|---|---|
| $H_2O$ | 1.28 | 1.15 | 1.20 | 1.10 | 1.14 | 1.17 |
| $D_2O$ | 1.11 | 0.93 | 1.02 | 1.10 | 1.13 | 1.06 |

**Table S23.** Raw data collected at 29.9 °C at pD 8.07 to evaluate isotope effect on $k_{cat}/K_M$ of thermolysin using 69 μM FAFLA.

**Table S24**

| Solvent | Approximated $k_{cat}/K_M$ ($\mu M^{-1}s^{-1}$) | | | | | Average ($\mu M^{-1}s^{-1}$) |
|---|---|---|---|---|---|---|
| $H_2O$ | 0.99 | 0.92 | 0.93 | 0.81 | 0.79 | 0.89 |
| $D_2O$ | 0.78 | 0.82 | 0.83 | 0.68 | 0.79 | 0.78 |

**Table S24.** Raw data collected at 17.3 °C at pD 8.07 to evaluate isotope effect on $k_{cat}/K_M$ of thermolysin using 69 μM FAFLA.

**Table S25**

| Phosphoramidon (mM) | $k_{app}$ ($s^{-1}$) | | |
|---|---|---|---|
| 0.000 | 0.002 | -0.005 | 0.003 |
| 0.103 | 0.115 | 0.112 | 0.115 |
| 0.051 | 0.074 | 0.074 | 0.065 |
| 0.077 | 0.097 | 0.100 | 0.103 |
| 0.026 | 0.040 | 0.043 | 0.044 |
| 0.013 | 0.022 | 0.026 | 0.023 |
| 0.256 | 0.222 | 0.217 | 0.255 |
| 0.205 | 0.214 | 0.205 | 0.210 |
| 0.513 | 0.304 | 0.310 | 0.294 |

**Table S25.** Raw saturation kinetics data collected at 29.9 °C at pH 8.00 of the binding of phosphoramidon to thermolysin.

**Table S26**

| 1/T ($K^{-1}$) | $k_{on}$ ($mM^{-1}s^{-1}$) | | |
|---|---|---|---|
| 0.003461 | 0.80 | 0.87 | 0.73 |
| 0.003412 | 0.98 | 0.93 | 0.92 |
| 0.003388 | 1.13 | 1.08 | 1.01 |
| 0.003366 | 1.26 | 1.16 | 1.08 |
| 0.003342 | 1.32 | 1.17 | 1.32 |
| 0.003321 | 1.48 | 1.39 | 1.30 |
| 0.003299 | 1.58 | 1.44 | 1.50 |
| 0.003278 | 1.67 | 1.77 | 1.53 |
| 0.003246 | 1.89 | 2.05 | 1.86 |

**Table S26.** Raw data of $k_{on}$ as a function of temperature at pH 8.00 using 17 µM phosphoramidon.

**Table S27**

| ZFLA (mM) | Rate ($s^{-1}$) | | |
|---|---|---|---|
| 0.00 | 35.8 | -28.6 | -7.2 |
| 0.14 | 217.1 | 206.1 | 96.1 |
| 0.28 | 291.3 | 213.8 | 238.3 |
| 0.43 | 301.7 | 398.4 | 398.6 |
| 0.57 | 462.7 | 452.1 | 472.3 |
| 0.71 | 584.9 | 509.5 | 384.5 |
| 1.06 | 496.5 | 529.1 | 668.1 |
| 1.42 | 648.1 | 653.1 | 714.2 |
| 2.83 | 781.5 | 905.3 | 763.2 |

**Table S27.** Raw Michaelis-Menten data collected at 29.9 °C at pH 7.20 using the ZFLA substrate.

**Table S28**

| **Condition** | **Break Point (°C)** | **$\Delta H°_c$ (kJ/mol)** | **$T\Delta S°_c$ (kJ/mol)** |
|---|---|---|---|
| Buffer | 28.0 | 34 ± 8 | 34 ± 8 |
| 1.1 M Ionic Strength | 24.0 | 28 ± 5 | 28 ± 5 |
| 2.5 M Ionic Strength | 26.4 | 18 ± 6 | 18 ± 6 |
| 10% Ficoll-70 | 25.4 | 18 ± 3 | 18 ± 3 |

**Table S28.** $\Delta H°_c$ and $T\Delta S°_c$ with physicochemical factors with break point temperature. Error is expressed as 95% confidence intervals.

**Table S29**

| **Condition** | **$k_{cat}$ ($s^{-1}$)** | **$K_M$ (mM)** |
|---|---|---|
| Buffer | 965 ± 63 | 0.55 ± 0.09 |
| 1.1 M Ionic Strength | 1370 ± 90 | 0.44 ± 0.08 |
| 5% Ficoll-70 | 866 ± 62 | 0.59 ± 0.10 |
| 10% Ficoll-70 | 701 ± 46 | 0.53 ± 0.09 |

**Table S29.** Steady state kinetic parameters of thermolysin with physicochemical factors at 29.9 °C and pH 7.20. Error is expressed as 95% confidence intervals.

**Table S30**

| pH | $k_{cat}$ ($s^{-1}$) | $K_M$ (mM) |
|---|---|---|
| 8.00 | 1360 ± 100 | 1.6 ± 0.2 |
| 7.50 | 1330 ± 70 | 1.1 ± 0.1 |
| 7.20 | 965 ± 63 | 0.55 ± 0.09 |

**Table S30.** Steady state kinetic parameters of thermolysin as a function of pH at 29.9 °C in the absence of ionic strength, viscosity, or macromolecular crowding. Error is expressed as 95% confidence intervals.

**Table S31**

| T (°C) | $k_{cat}$ ($s^{-1}$) | $K_M$ (mM) |
|---|---|---|
| 29.9 | 1360 ± 100 | 1.6 ± 0.2 |
| 23.8 | 1090 ± 80 | 1.5 ± 0.2 |
| 17.3 | 1000 ± 50 | 1.7 ± 0.2 |

**Table S31.** Steady state kinetic parameters of thermolysin as a function of temperature at pH 8.00 in the absence of ionic strength, viscosity, or macromolecular crowding. Error is expressed as 95% confidence intervals.

**Table S32**

| [Sucrose] (w/w) | $k_{cat}$ ($s^{-1}$) | $K_M$ (mM) |
|---|---|---|
| 0 | 1360 ± 100 | 1.6 ± 0.2 |
| 10 | 1230 ± 80 | 1.2 ± 0.2 |
| 20 | 835 ± 39 | 0.80 ± 0.09 |
| 25 | 642 ± 33 | 0.77 ± 0.09 |
| 30 | 546 ± 22 | 0.65 ± 0.07 |

**Table S32.** Steady state kinetic parameters of thermolysin with elevated viscosity at 29.9 °C and pH 8.00. Error is expressed as 95% confidence intervals.

**Table S33**

| [Sucrose] (w/w) | $k_{cat}$ ($s^{-1}$) | $K_M$ (mM) |
|---|---|---|
| 0 | 1000 ± 50 | 1.7 ± 0.2 |
| 20 | 564 ± 26 | 0.73 ± 0.09 |

**Table S33.** Steady state kinetic parameters of thermolysin with elevated viscosity at 17.3 °C and pH 8.00. Error is expressed as 95% confidence intervals.

**Table S34**

| $D_2O$ (%) | $k_{cat}$ ($s^{-1}$) | $K_M$ (mM) |
|---|---|---|
| 0 | 1330 ± 70 | 1.1 ± 0.1 |
| 20 | 1130 ± 40 | 0.86 ± 0.08 |
| 33 | 1080 ± 70 | 0.85 ± 0.13 |
| 60 | 952 ± 49 | 0.76 ± 0.09 |

**Table S34.** Steady state kinetic parameters of thermolysin with elevated $D_2O$ at 29.9 °C and pH 7.50 and pD 8.07. Error is expressed as 95% confidence intervals.

**Figure S1**

A

B

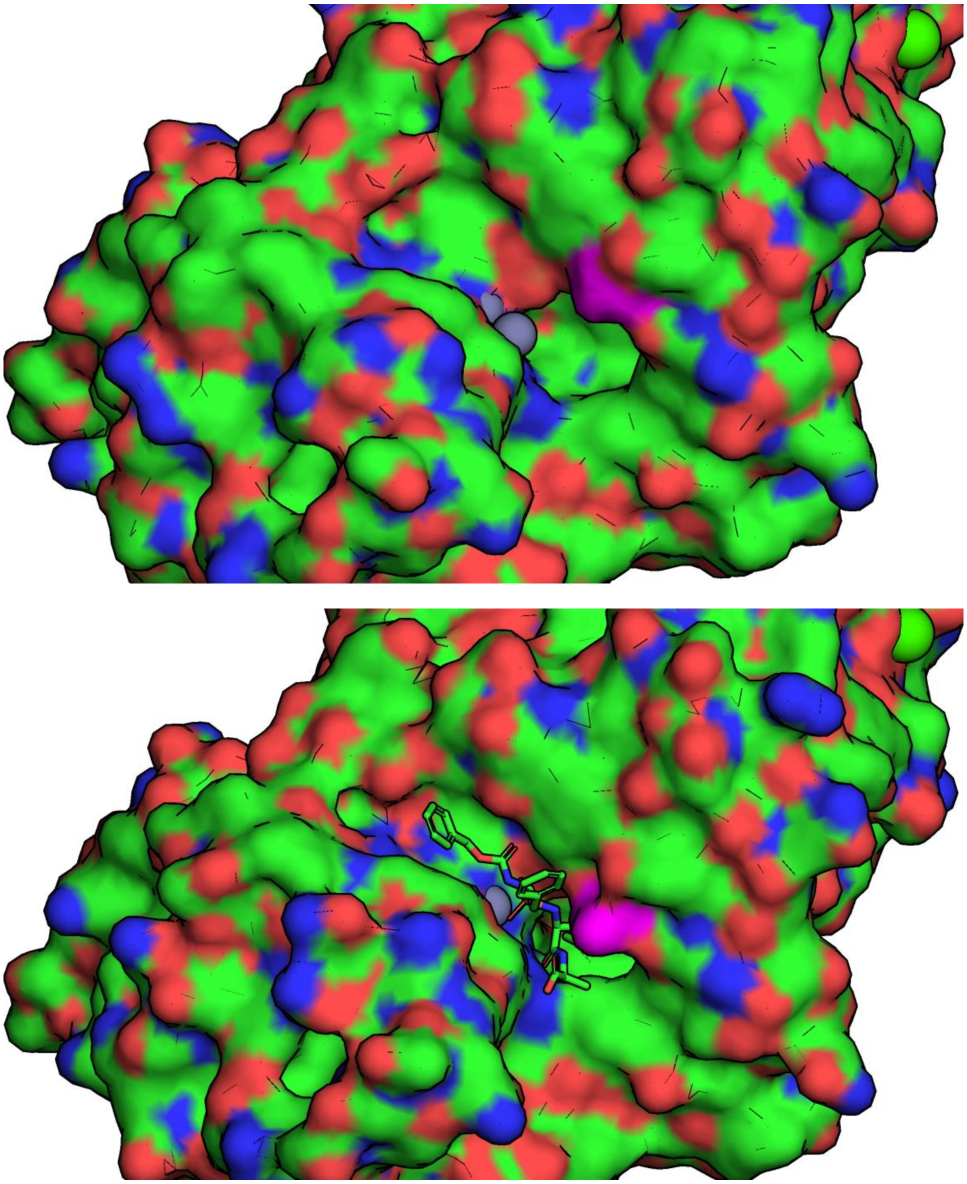

**Figure S1.** Hinge Bending Motions of Thermolysin. A. Open form thermolysin (PDB 1l3f)[28], with excess zinc bound, representative of the ligand-free enzyme. B. Closed form thermolysin (PDB 4tmn)[24], with $ZF^{P}LA$ bound, representative of the ligand-bound enzyme. Asn112, which is a major residue involved in this conformational change, is shown in purple for both structures.

**Figure S2**

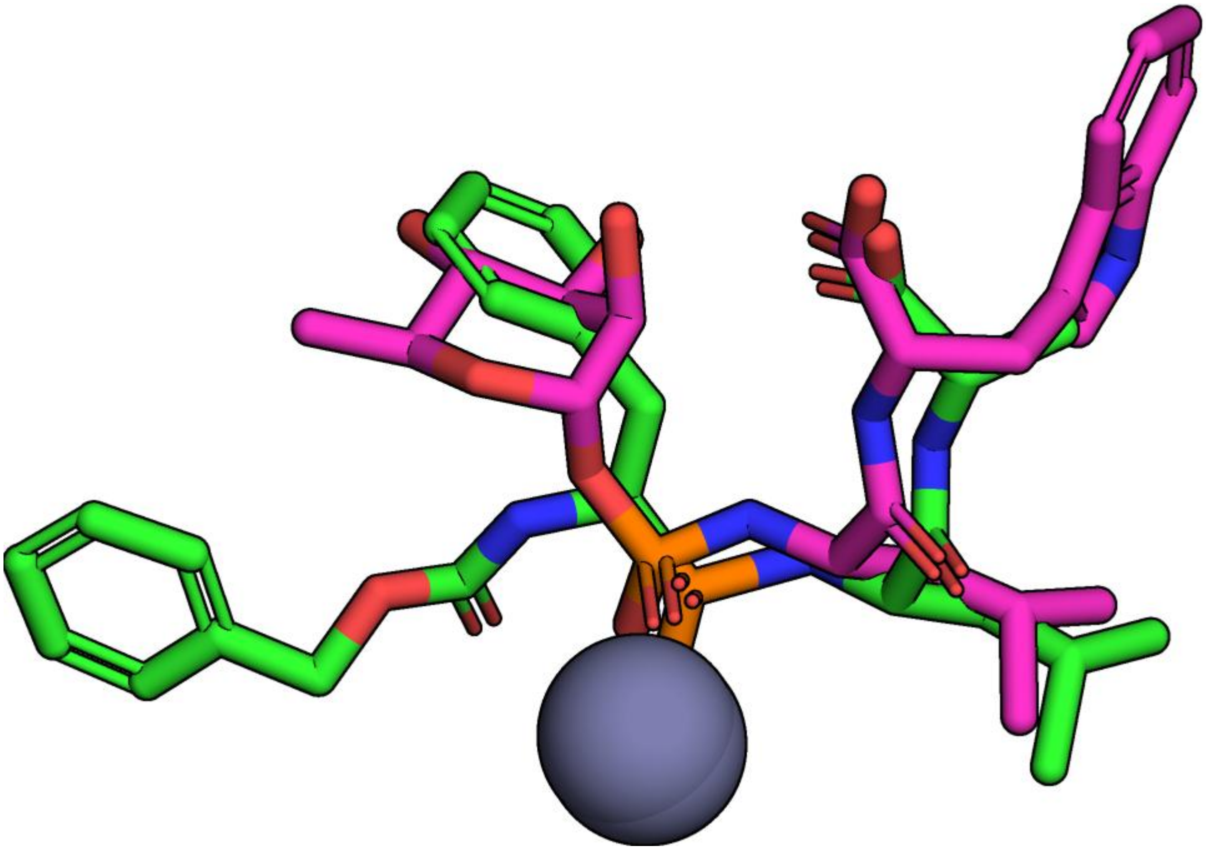

**Figure S2.** Binding of ZF$^P$LA and phosphoramidon to thermolysin. ZF$^P$LA (green from PDB 4tmn)[24], the transition state analog of FAFLA, binds in a similar mode to phosphoramidon (magenta from PDB 1TLP)[29]. Of particular interest is the deoxymannose ring of phosphoramidon being in a similar position to the Phe residue of ZF$^P$LA, which may preserve some of the steric interactions with Asn112 which occur for FLA ligands like FAFLA.

**Figure S3**

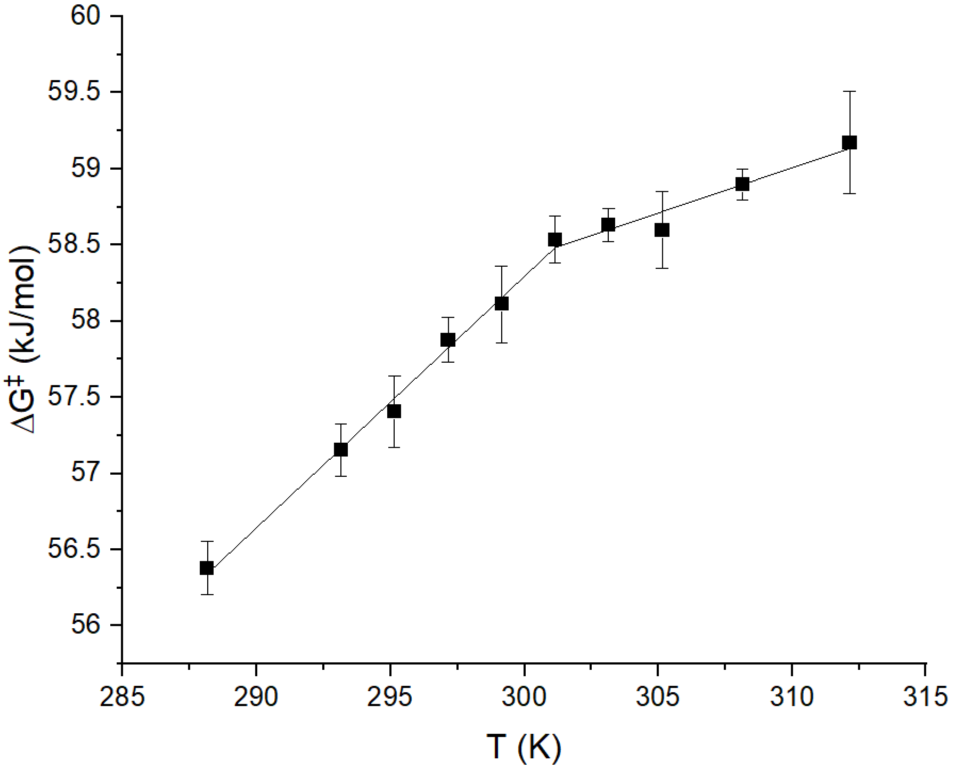


**Figure S3.** Temperature Dependence of ΔG‡ for Thermolysin with the FAFLA Substrate. ΔG‡ shows a distinct break point at 28 °C after which the rate of increase as a function of temperature decreases. Similar to succinylcasein, the increase in ΔG‡ with temperature indicates that the rate increase observed with temperature is due to increased thermal energy rather than a decreased activation barrier. Error bars represent 95% confidence intervals.

**Figure S4**

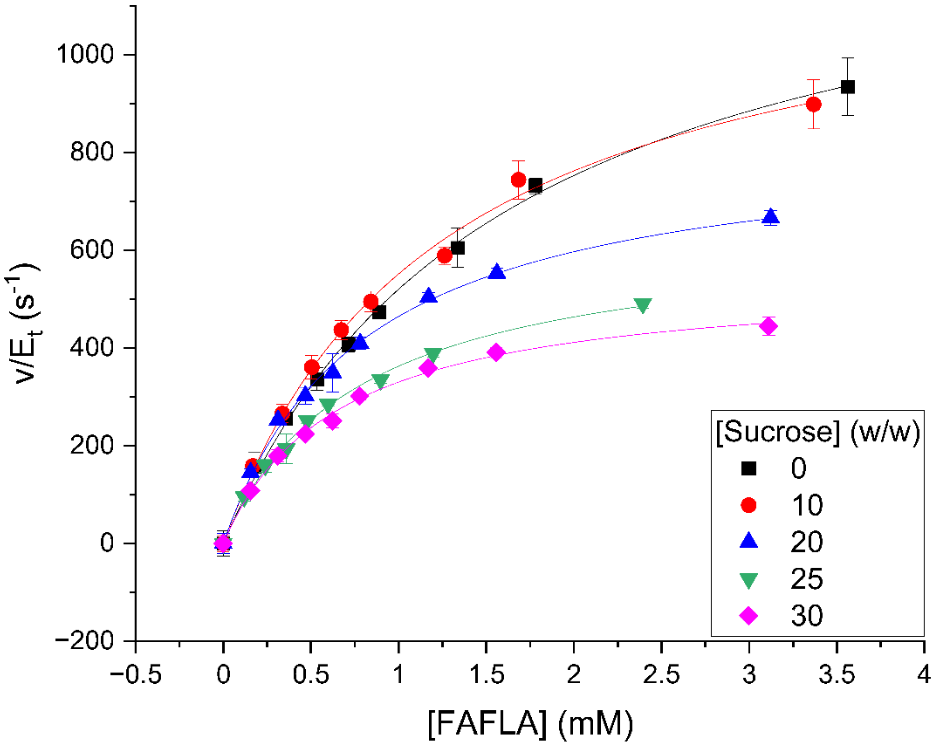


**Figure S4.** Michaelis-Menten Plots of Thermolysin with Elevated Viscosity. Both the $k_{cat}$ and $K_M$ for FAFLA of thermolysin decreased with elevated viscosity caused by the addition of sucrose. The $k_{cat}$ decrease enabled KSVEs to be evaluated and it was found that this parameter is almost fully dependent on viscosity. Experiments were performed at pH 8.00. Error bars represent 95% confidence intervals. Table S32 contains Michaelis-Menten parameters obtained from these fits.

**Figure S5**

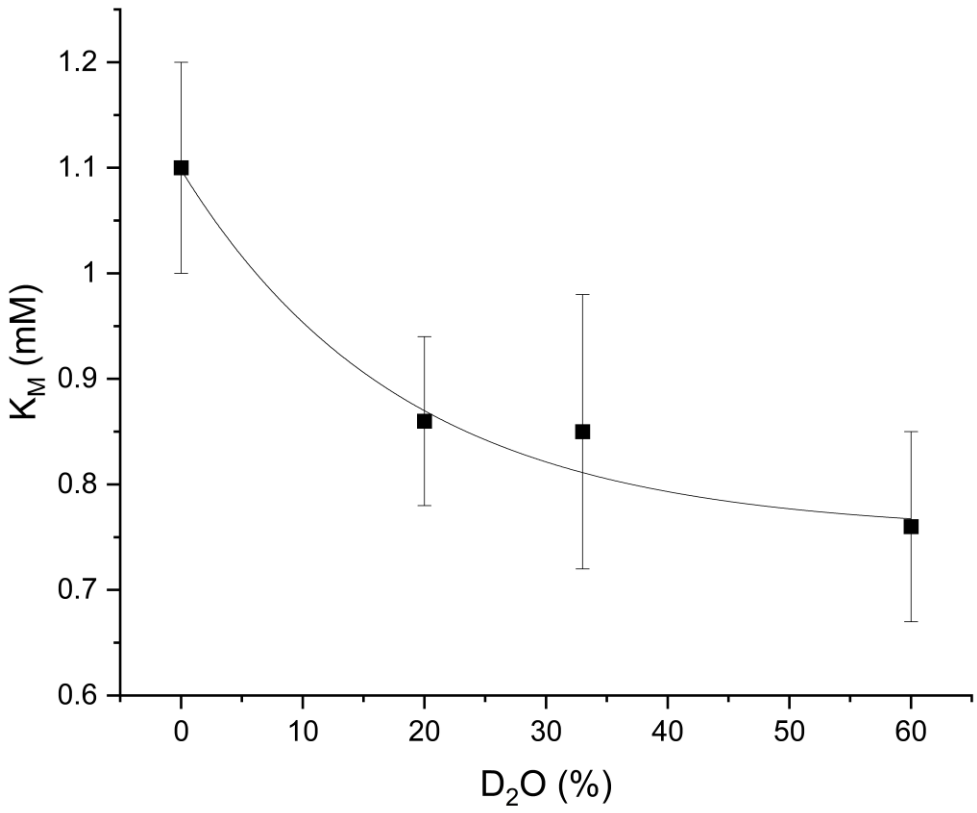


**Figure S5.** Effect of Increasing Concentrations of $D_2O$ on the $K_M$ for FAFLA of Thermolysin. An exponential decay fit to the data indicates a $K_M$ of 0.75 mM (1.45-fold decrease) at 100% $D_2O$, using the equation $y = 0.75 + 0.34e^{-x/18.3}$. The experiments were performed at pH 7.50. Error bars represent 95% confidence intervals. Table S34 contains values used to construct this figure.

### Figure S6

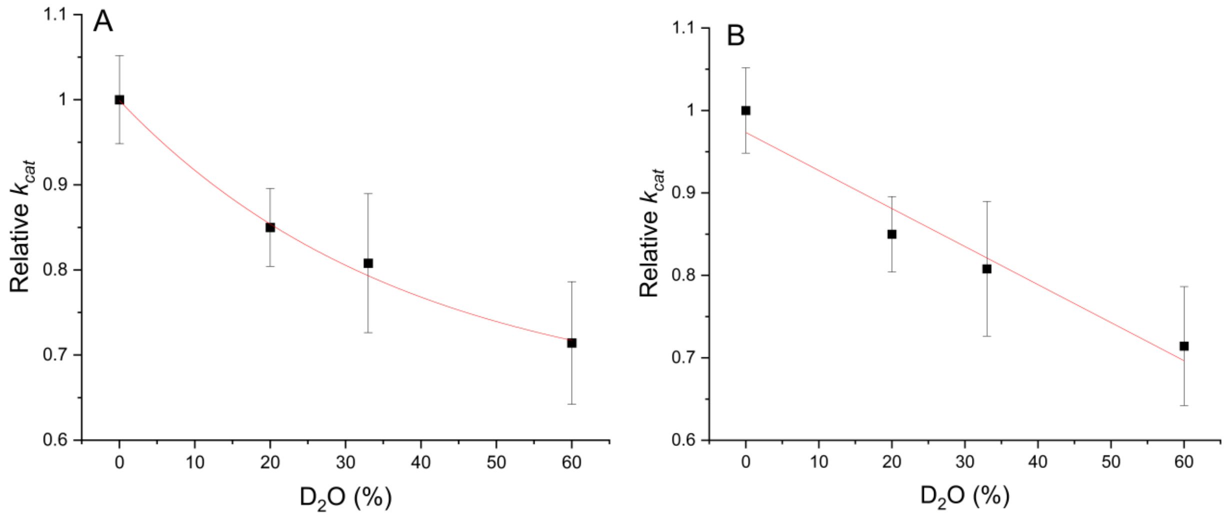


**Figure S6.** Curve Fitting of the Proton Inventory of Thermolysin. A. Proton inventory fit with an exponential decay fit predicts a 1.5-fold decrease in $k_{cat}$ at 100% $D_2O$ with an $R^2$ of 0.99. B. Proton inventory fit with a linear fit predicts a 2.0-fold decrease in $k_{cat}$ at 100% $D_2O$ with an $R^2$ of 0.92. Only the exponential decay model, coupled with the $K_M$ data in Figure S5, provides a prediction of $k_{cat}/K_M$ at 100% $D_2O$ which is consistent with experimental results in Figure 3B. Error bars represent 95% confidence intervals. Table S34 contains values used to construct this figure.

### Figure S7

The $k_{cat}$ of thermolysin with the FAFLA substrate is 91% sensitive to viscosity and 100% $D_2O$ is 22% more viscous than $H_2O$ at 29.9 °C[34]. This means that 100% $D_2O$ should exert a 1.20-fold decrease (91% of 1.22 relative viscosity) on the $k_{cat}$ due to only viscosity. The $k_{cat}$ would decrease from 1330 $s^{-1}$ in 100% $H_2O$ to 1110 $s^{-1}$ in 100% $D_2O$ due to this viscosity effect.

The $k_{cat}$ of thermolysin is predicted by the exponential fit in Figure S6 to decrease by 1.50-fold at 100% $D_2O$, resulting in a value of 888 $s^{-1}$. This experimentally-determined value is 1.25-fold larger than the effect predicted by viscosity, thus the contribution of the presence of deuterium is a KSIE on the $k_{cat}$ of 1.25.

**Figure S7.** Deconvolution of Viscosity Effects from Isotope Effects. By considering the effect of increased viscosity of $D_2O$, the effect of the presence of deuterium alone can be elucidated.

**Figure S8**

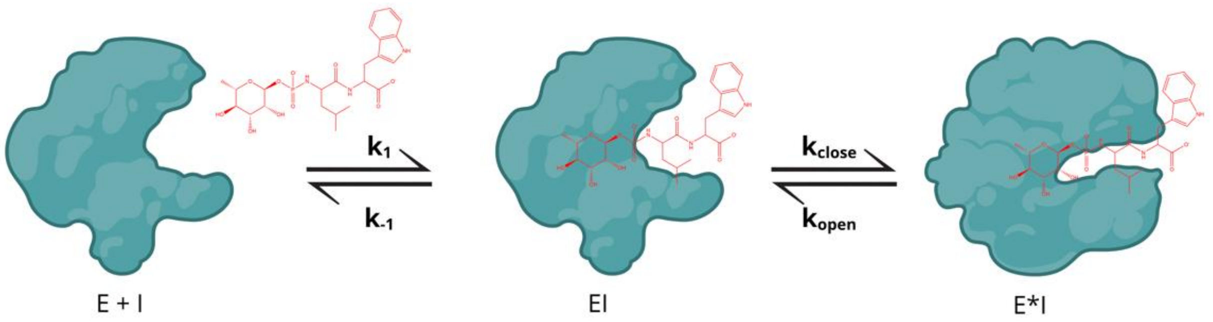


**Figure S8.** Kinetic Scheme for the Binding of Phosphoramidon to Thermolysin. Thermolysin first binds phosphoramidon in the open form which is modeled by the bimolecular association rate constant $k_1$ [35-36].The release of phosphoramidon from the open form is modeled by $k_{-1}$. Upon ligand binding, thermolysin undergoes a slow, unimolecular open-to-closed conformational change, which is described by the rate constant $k_{close}$ [35-36]. Desolvation of the active site during this closing step causes the fluorescence intensity of Trp115 to increase[35-38]. The reverse closed-to-open conformational change, which is analogous to product release, is described by $k_{open}$.

### Figure S9

Fluorescence titrations across a temperature range of 16-35 °C were performed to evaluate the temperature dependence of the $K_d$ for phosphoramidon with an Agilent Eclipse fluorometer[35-38]. A thermolysin concentration of 1.14 μM was used for all temperatures except for 35 °C, where 0.930 μM was used. Concentrations of phosphoramidon were determined using $\varepsilon_{280}$ of 5.55 $mM^{-1}cm^{-1}$ and ranged from 0-2.5 μM. The experiments were performed in 50 mM sodium borate buffer (pH 8.00). The samples were incubated at room temperature for 30 min to allow the system to reach equilibrium. Samples were equilibrated to the desired temperature for 3 min in an external water bath and quickly transferred to the fluorometer for measurements. The excitation wavelength was set to 280 nm and emission was collected at 360 nm. Triplicate data was collected using three independently prepared samples of thermolysin mixed with phosphoramidon. Additionally, fluorescence intensity of free phosphoramidon and thermolysin at the same concentrations used in the titration were recorded for use with Eq. S1.

The data is fit against inhibitor concentration as values of $\Delta f$ (%) according to Eq. S1. The numerator refers to the difference in fluorescence intensity of the EI complex compared to the sum of free E and free I, and the denominator is the sum of fluorescence intensity of enzyme and an equivalent concentration of phosphoramidon. Since phosphoramidon has been shown to bind to thermolysin with nM affinity such that $K_d \ll [E]$, the data was fit to Morrison's quadratic equation for tight binding inhibitors in Origin v.2024 software (Eq. S2)[35-38], where $\Delta F_{max}$ and $K_d$ were determined by the fit and $E_t$ was given as a fixed value.

Equation S1. $$\Delta f\ (\%) = \frac{F_{EI} - (F_E + F_I)}{F_E + F_{I,eq}}$$

Equation S2. $$\Delta f\ (\%) = \Delta F_{max} \frac{E_t + I + K_d - \sqrt{(E_t + I + K_d)^2 - 4E_t I}}{2E_t}$$

**Figure S9.** Fluorescence Titration to Measure Kd of Thermolysin and Phosphoramidon. Procedure for fluorescence titration of thermolysin with the phosphoramidon inhibitor was adapted from Kitagishi and Hiromi [35-38].

**Figure S10**

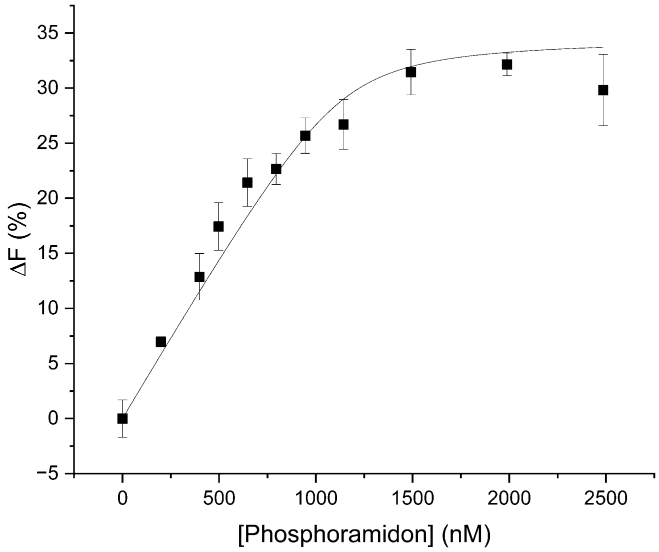


**Figure S10.** Example Fluorescence Titration of Thermolysin with Phosphoramidon. Thermolysin (1.14 µM) was titrated with varying concentrations of phosphoramidon according to Kitagishi and Hiromi[35-38]. The resulting $K_d$ values have unacceptably large error since the $K_d$ is 20-fold lower than enzyme concentration used.

**Figure S11**

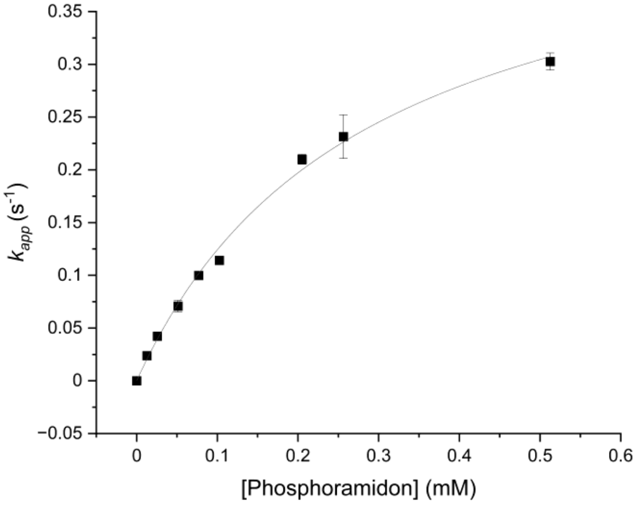


**Figure S11.** Hyperbolic Plot of $k_{app}$ as a Function of Phosphoramidon Concentration. The above plot shows that binding of phosphoramidon to thermolysin follows saturation kinetics, which is due to the interplay of a fast bimolecular binding step followed by a slow unimolecular hinge bending step[35-36]. Values of $k_{close}$ and $K_{sc}$ at pH 8.00 were 0.48 ± 0.04 $s^{-1}$ and 0.28 ± 0.05 mM, respectively, and directly obtained from the fit. Error bars represent 95% confidence intervals.

**Figure S12**

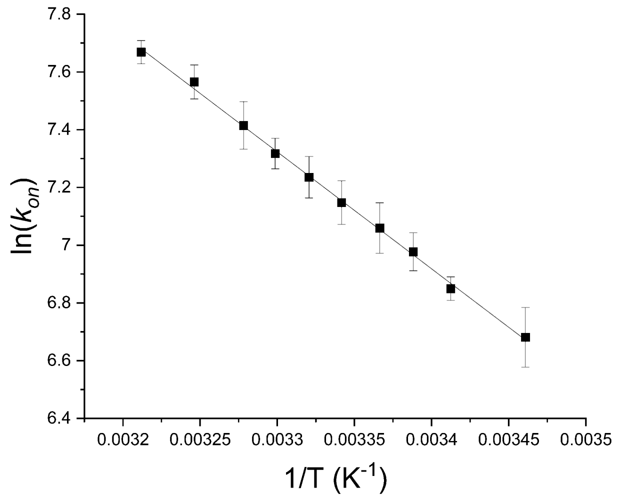


**Figure S12.** Temperature Dependence of Hinge Bending Motions of Thermolysin. $k_{on}$, represented by $\frac{k_{close}}{K_{SC}}$, shows monophasic behavior across the full temperature range. Since the $K_d$ for phosphoramidon binding is likely linear with temperature like other inhibitors[17,38], $k_{off}$ ($k_{open}$) should be linear as well. This indicates that the hinge bending motions during the catalytic cycle are not the rate limiting step responsible for concave break point shape. Error bars represent 95% confidence intervals.

**Figure S13**

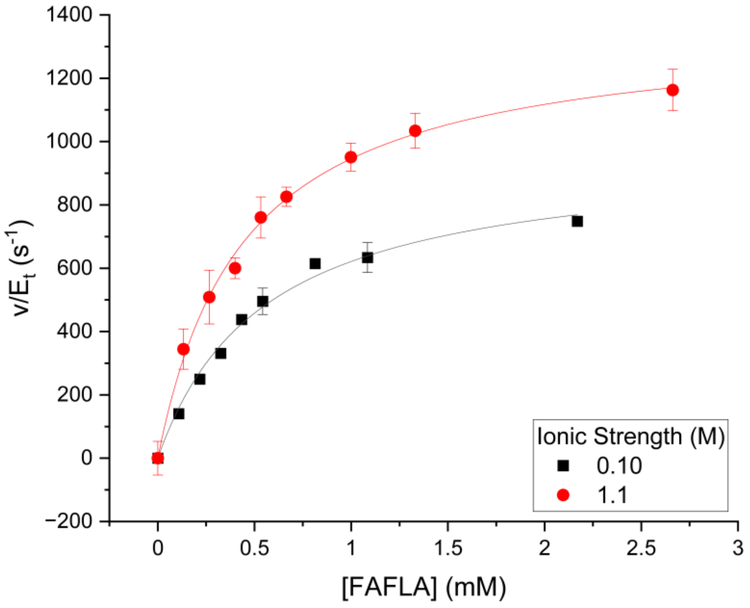


**Figure S13.** Michaelis-Menten Kinetics of Thermolysin in Elevated Ionic Strength. The $k_{cat}$ of thermolysin increased 1.40-fold between 0.1-1.1 M total ionic strength with no significant change to $K_M$ for FAFLA. Experiments performed at pH 7.20. Error bars represent 95% confidence intervals. Table S29 contains kinetic parameters obtained from these fits.

**Figure S14**

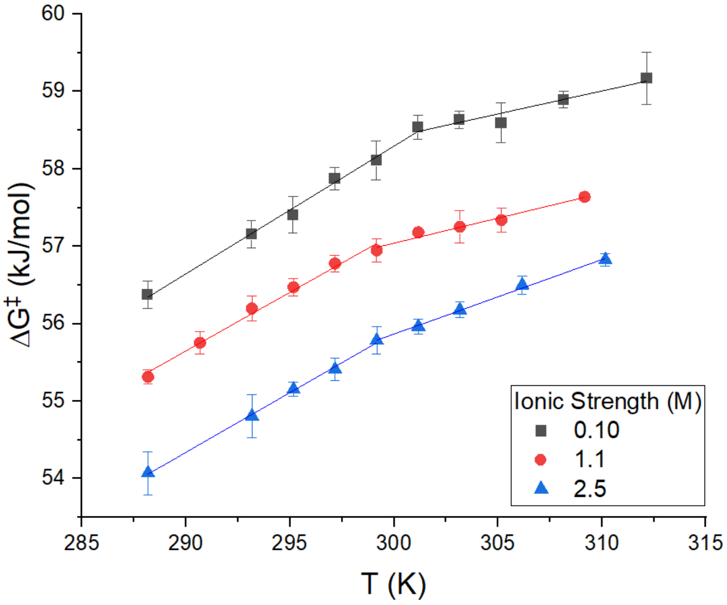


**Figure S14.** Temperature Dependence of ΔG‡ with Elevated Ionic Strength. ΔG‡ becomes increasingly linear between ionic strengths of 0.10-2.5 M total ionic strength, mirroring results with the succinylcasein substrate. Experiments performed at pH 7.20. Error bars represent 95% confidence intervals.

---

**Figure S15**

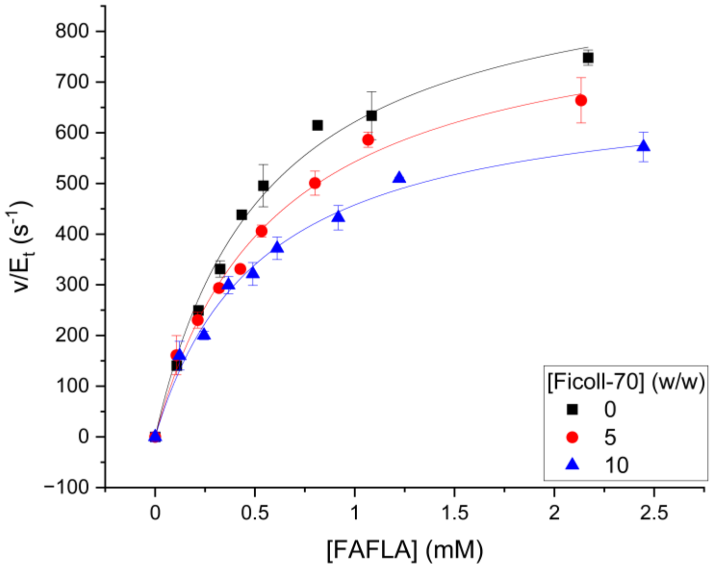


**Figure S15.** Michaelis-Menten Kinetics of Thermolysin in Elevated Macromolecular Crowding. The $k_{cat}$ of thermolysin decreased 1.37-fold between 0-10% Ficoll-70 with no significant change to $K_M$ for FAFLA. Only one trend was observed with the FAFLA substrate, unlike the biphasic effects of crowding with the succinylcasein substrate. Experiments performed at pH 7.20. Error bars represent 95% confidence intervals. Table S29 contains kinetic parameters obtained from these fits.

---

**Figure S16**

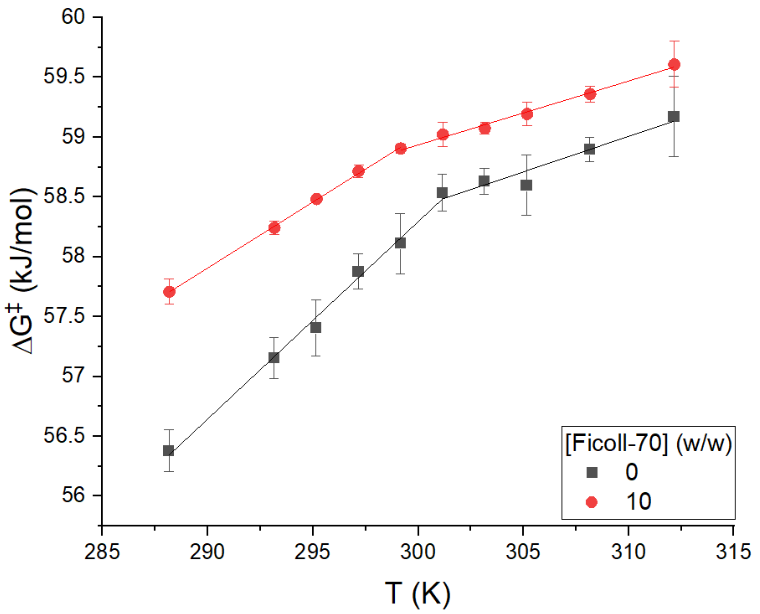


**Figure S16.** Temperature Dependence of $\Delta G^{\ddagger}$ with Elevated Macromolecular Crowding. $\Delta G^{\ddagger}$ becomes increasingly linear between 0-10% Ficoll-70, mirroring results with the succinylcasein substrate. Experiments performed at pH 7.20. Error bars represent 95% confidence intervals.

---

**Figure S17**

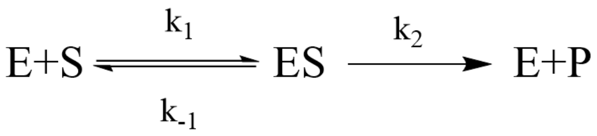


The standard Michaelis-Menten model (above) derives the kinetic parameters as follows, such that the rate constants that describe the initial bimolecular binding step are not represented within $k_{cat}$.

$$k_{cat} = k_2 \text{ and } \mathrm{K_M} = \frac{k_{-1} + k_2}{k_1}$$

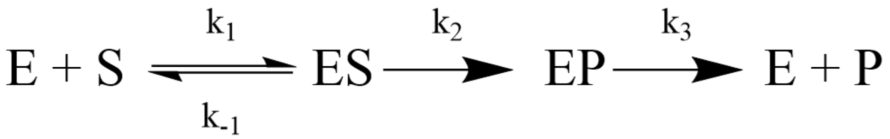


The same is true of an expanded Michaelis-Menten expression that treats product release as a distinct step[31].

$$k_{cat} = \frac{k_2 k_3}{k_2 + k_3} \text{ and } \mathrm{K_M} = \frac{k_{-1} + k_2}{k_1} x \frac{k_3}{k_2 + k_3}$$

**Figure S17.** Bimolecular Binding Rate Constants do not Influence $k_{cat}$. Only unimolecular, saturable processes are represented within $k_{cat}$, therefore the initial bimolecular binding is not responsible for concavity.

---

**Figure S18**

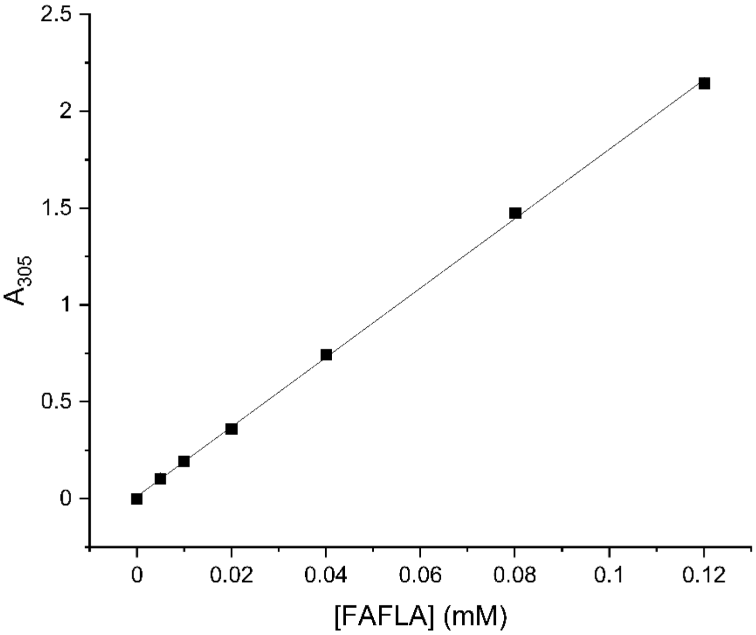


**Figure S18.** Extinction Coefficient of FAFLA at 305 nm. The extinction coefficient of the full length FAFLA substrate was determined to be 17.9 ± 0.4 $mM^{-1}cm^{-1}$ which was used to determine stock concentration of substrate. Error bars represent 95% confidence intervals.

---

**Figure S19**

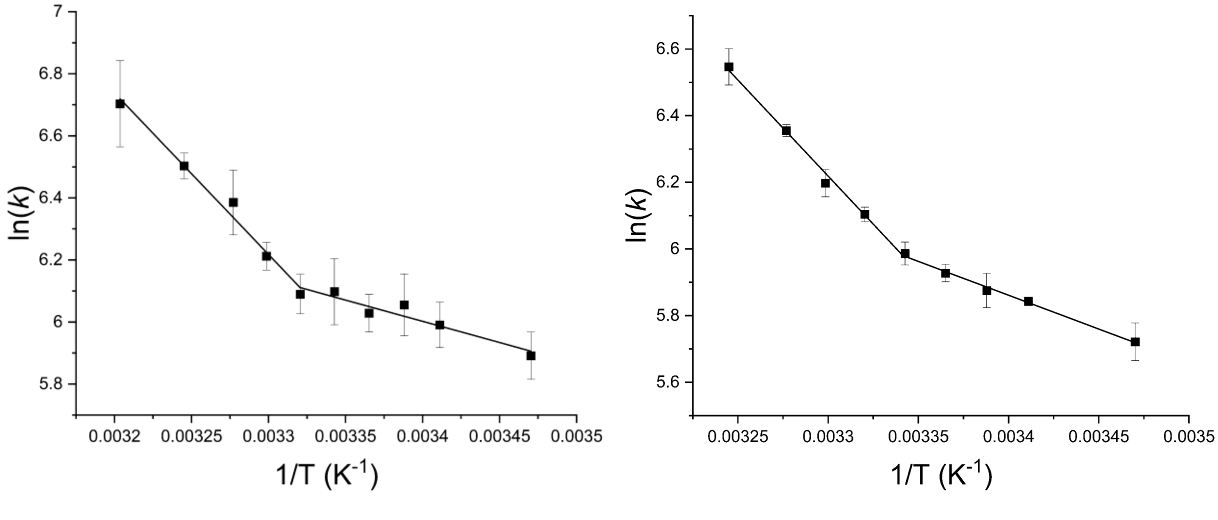


**Figure S19.** Thermolysin Arrhenius Plots at pH 7.20 and 8.00. There is no difference in $\Delta H°_c$ and $T\Delta S°_c$ between pH 7.20 (A) (34 ± 8 kJ/mol) and pH 8.00 (B) (31 ± 6 kJ/mol). Error bars represent 95% confidence intervals.

---

**Figure S20**

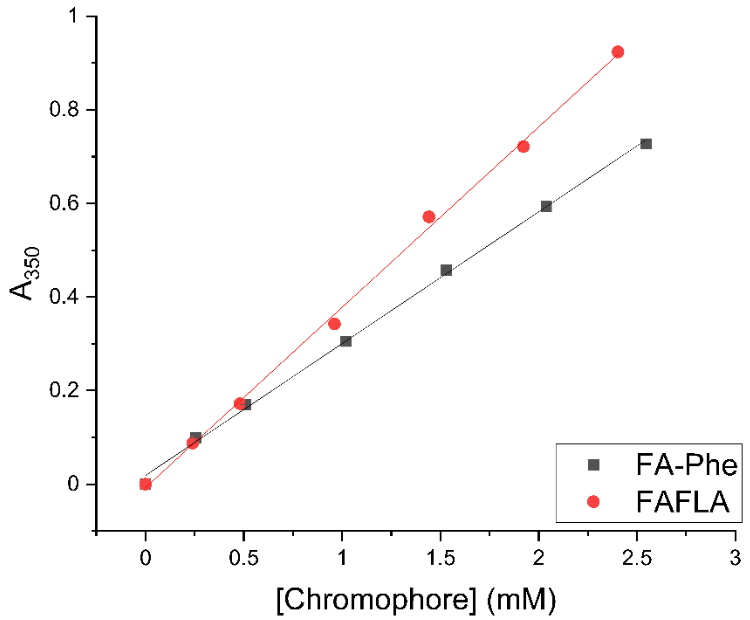


**Figure S20.** Difference Extinction Coefficient Between FAFLA and FA-Phe at 350 nm. The difference extinction coefficient between full length FAFLA and the FA-Phe product was determined to be -0.104 ± 0.020 $mM^{-1}s^{-1}$, which is the extinction coefficient used to monitor thermolysin-catalyzed cleavage of FAFLA. 350 nm was chosen because the absorbance of FAFLA is sufficiently low at concentrations above $K_M$ (~1-1.5 mM) to enable viable determinations of $k_{cat}$ and $K_M$. Error bars represent 95% confidence intervals.

---

**Figure S21**

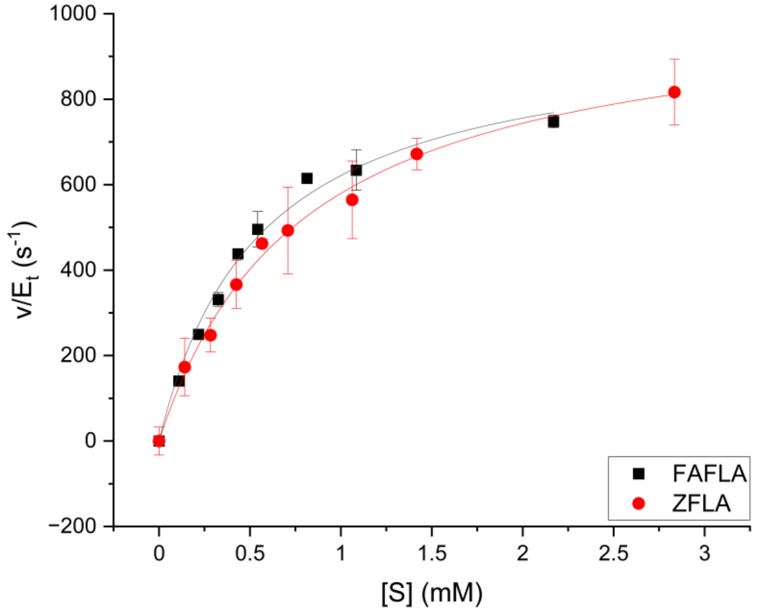


**Figure S21.** Verification of Difference Extinction Coefficient at 350 nm. The $k_{cat}$ and $K_M$ for FAFLA of thermolysin were found to be 960 ± 60 $s^{-1}$ and 0.55 ± 0.09 mM, respectively. The $k_{cat}$ and $K_M$ for ZFLA of thermolysin were found to be 1000 ± 130 $s^{-1}$ and 0.79 ± 0.23 mM, respectively. Both sets of values are agreeable with the literature[20,21]. The trinitrobenzenesulfonic acid-linked assay previously described[1] was used to monitor ZFLA cleavage and ZFLA was synthesized by Biomatik. The similarity between kinetic parameters for the two nearly identical peptides indicated that the extinction coefficient in Figure S21 is reflective of cleavage of FAFLA into FA-Phe. Experiments performed at pH 7.20 and 29.9 °C. Error bars represent 95% confidence intervals.

**Figure S22**

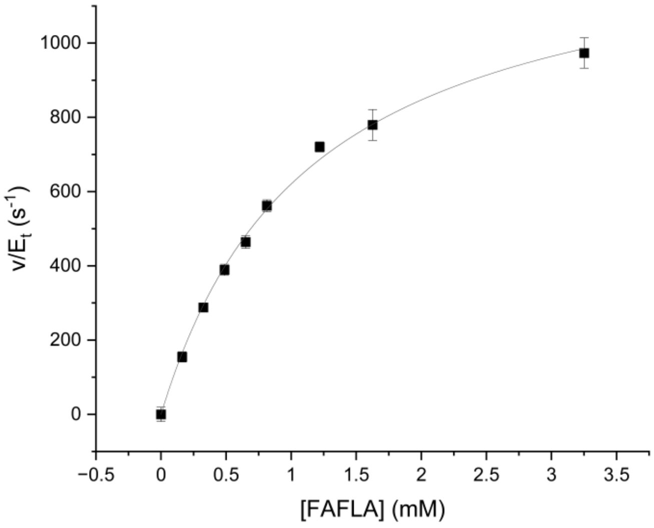


**Figure S22.** Kinetic Parameters of Thermolysin at pH 7.50. The $k_{cat}$ and $K_M$ of thermolysin at pH 7.50 were found to be 1330 ± 70 $s^{-1}$ and 1.1 ± 0.1 mM, respectively. The pH had to be adjusted for KSIE experiments to enable comparison with the literature[19,32]. These kinetic parameters enabled a proper FAFLA concentration (≤10-fold less than $K_M$) to be used to approximate values of $k_{cat}/K_M$ at 100% $D_2O$. Error bars represent 95% confidence intervals.

**Figure S23**

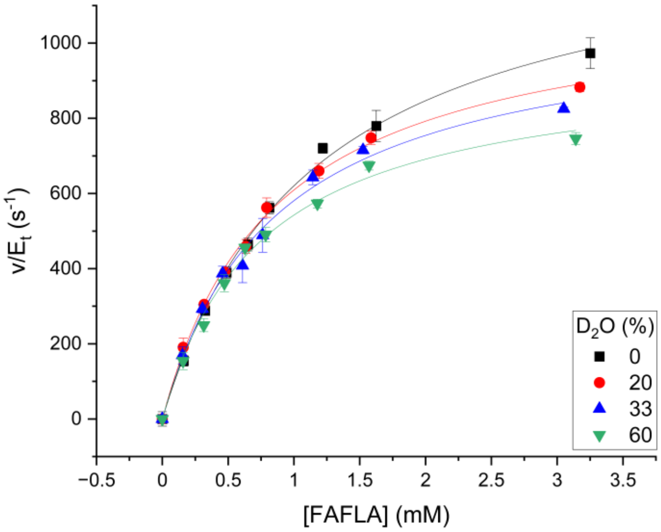


**Figure S23.** Michaelis-Menten Plots of Thermolysin with Increasing $D_2O$. Both the $k_{cat}$ and $K_M$ for FAFLA of thermolysin decreased with increasing concentrations of $D_2O$, enabling the KSIE on the $k_{cat}$ to be elucidated. Experiments performed using mixtures of pH 7.50 and pD 8.07 buffer. Error bars represent 95% confidence intervals. Table S34 contains kinetic parameters obtained from these fits.

**Figure S24**

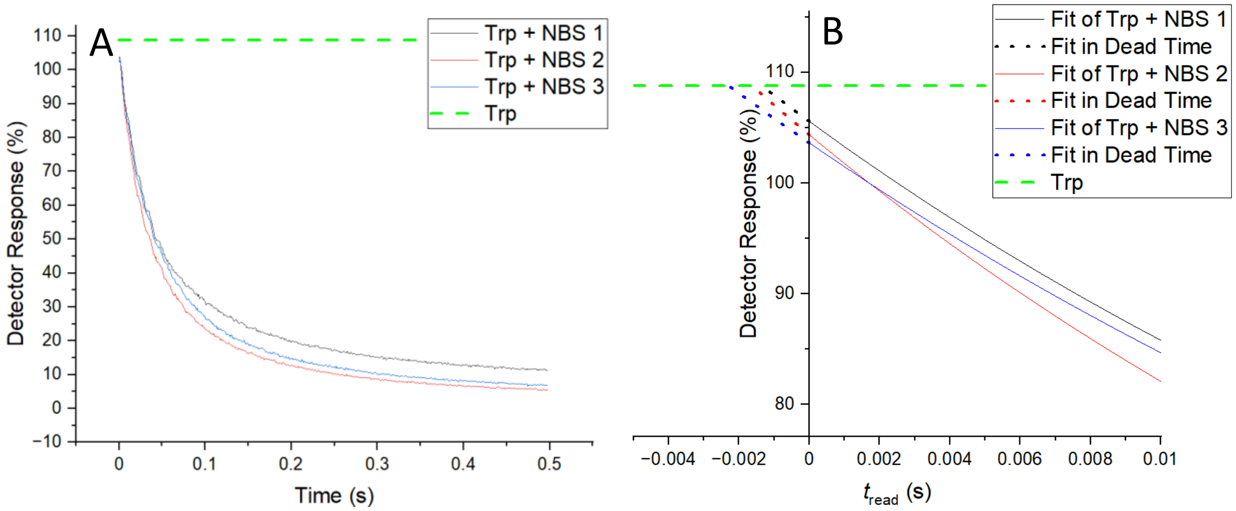


**Figure S24.** Stopped Flow Dead Time Calculation. A. Tryptophan (Trp) at 10 µM and N-bromosuccinimide (NBS) at 100 µM each dissolved in 50 mM sodium borate buffer (pH 8.00) were used to determine the dead time of the stopped flow instrument. The excitation wavelength was set to 280 nm, fluorescence quenching over the 0.5 s time course was recorded in triplicate, and fit to an exponential decay model. B. Extrapolation of the fit into negative time to reach the fluorescence of Trp alone (green dashed line) enabled a dead time of 1.8 ± 0.5 ms to be determined.

**Figure S25**

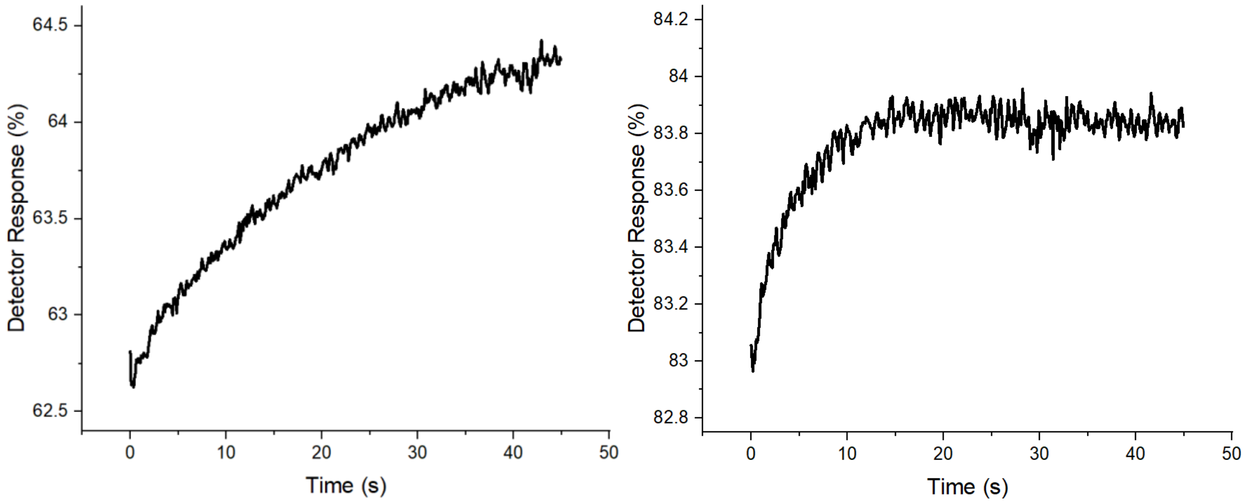


**Figure S25.** Example Time Courses of Phosphoramidon Binding to Thermolysin. The time courses are representative of the increase in fluorescence vs time due to inhibitor binding to thermolysin and formation of the mature closed complex at 0.026 mM (A) and 0.256 mM (B) phosphoramidon at pH 8.00. Rate constants obtained from these fits were used to construct Figure S11.
